# Cancer cells hijack mitochondria from Bhlhe40+ T cells for immune evasion

**DOI:** 10.64898/2026.10.23.761277

**Authors:** Hailong Hu, Yuhao Tan, Irina Shkundina, Richard Madnick, Jing Hu, Hongyi Zhang, Ziqi Yang, Tong Wang, Xuexin Yu, Julia Fleming, Mingyao Pan, Yizhou Wang, Yuanyuan Pang, Drew Weissman, Bo Li

## Abstract

Cancer cells hijack mitochondria from T cells via tunneling nanotubes (TNTs) to promote immune evasion, yet the underlying mechanisms of mitochondrial-loss remain poorly understood. Here, through targeted screening of relevant genes, we demonstrated that CD38 is a potent regulator of TNT formation through interaction with its cognate ligand, CD31. Conditional knockdown of Cd38 in murine models leads to CD8^+^ T cell-dependent tumor shrinkage. Single-cell RNA sequencing of tumor-infiltrating T cells revealed that cancer cells preferentially target the Bhlhe40+ effector subset. Bhlhe40 is a transcription factor that upregulates the expression of S100a4, a reported chemoattractant of nanotubes. Bhlhe40 upregulation creates mitochondrial-loss vulnerability for tumor neoantigen-specific T cells. Dual mitochondria transfer blockade with Cd38 knockdown and S100a4 inhibitor trifluoperazine markedly suppressed tumor growth. Collectively, our findings identified the Bhlhe40–S100a4 cascade as a critical regulator of cancer/T cell mitochondrial transfer, providing insights for the development of new cancer immunotherapies.

## Introduction

In the past two decades, many studies have collectively established that mitochondrion, an essential intracellular organelle for energy production, can be horizontally transferred between mammalian cells^1,2^. This process is predominantly mediated by tunneling nanotubes, or TNTs, which are open-ended membrane protrusions that directly connect the cytosol of distal cells^2,3^. A variety of cell types can be connected via TNTs to transfer mitochondria as donors or acceptors, including neurons, astrocytes, heart muscle cells, epithelial cells, stem cells and immune cells^1–6^. This instant mitochondrial transfer serves as a quick alleviation of oxidative stress to mitigate severe tissue damage^7,8^.

Using imaging and genetic approaches, we and others have demonstrated that a variety of cancer cells may exchange mitochondria with T cells^9–12^. During this process, cancer cells either unidirectionally hijack T cell mitochondria^9,11^ or transfer defective mitochondria back, thereby promoting T cell metabolic dysfunction^10,12^. Non-specific disruption of TNT formation via farnesylation inhibitors reduced mitochondrial transfer *in vitro* and *in vivo*, and has shown synergy with immune checkpoint blockade therapies^11^. Further, as the primary mediators of anti-tumor cytotoxicity, tumor-reactive T cells may be particularly susceptible to mitochondrial loss due to sustained and intimate physical interactions with cancer cells, providing a potential mechanistic pathway for immune evasion.

Despite these exciting insights, critical knowledge gaps remain. First, T cells are not typical mitochondrial donors, and thus it is unclear how cancer cells target T cells for mitochondrial hijacking. In addition, multiple T-cell subsets coexist in the tumor microenvironment, such as cytotoxic CD8^+^ T cells, conventional helper CD4^+^ T cells and regulatory T cells. Immune evasion requires cancer cells to selectively target CD8^+^ or helper CD4^+^ T cells, while keeping T-regs intact. However, the underlying mechanism for target differentiation is completely unknown. Finally, the behavior of (neo)antigen-specific T cells during mitochondrial transfer remains poorly characterized. It is unclear if their closer physical distance to cancer cells will lead to higher mitochondrial exchange activities, and ultimately, dysfunctional anti-tumor immune responses. Clearly, addressing these questions will advance the understanding of cancer-T cell interaction and create novel therapeutic opportunities for cancer immunotherapies.

Here, we filled these gaps by controlling nanotube formation without cytoskeleton disruption to characterize how mitochondrial transfer reshapes T cell phenotypes *in vivo*. Through unbiased high-throughput genomic assay, we identified that Basic Helix Loop-Helix Family Member e40 (BHLHE40), a transcription factor that regulates T cell effector function, plays a central role in mitochondrial-hijacking vulnerability. Mechanistically, BHLHE40 upregulates S100 calcium-binding protein A4 (S100A4), which is a chemoattractant for nanotube guidance. Higher expression of BHLHE40 creates multiple channels for mitochondrial transfer between cancer and effector T cells, leading to elevated ROS production and accelerated T cell apoptosis *in vivo*. Blockade of this pathway protected neoantigen-specific T cell functionality and significantly reduced tumor growth. Together, this study revealed a previously unreported mechanism that governs T cell mitochondrial loss in the tumor microenvironment.

## Results

### Target screen for intercellular mitochondrial transfer blockade

To establish an *in vitro* system for mitochondria transfer, we tested 3 human (A549, U2OS, PC9) and 2 murine (MC38 and B16/F10) cell lines to perform coculture with primary T cells (**Figure S1A**). We first stained mitochondria of A549 cells with MitoTracker^Green^ and primary CD8+ T cell with MitoTracker^DeepRed^ (**Figures 1A and S1B**). A549 and T cells were mixed at 5:1 ratio cocultured for 3 hours, and separated by flow sorting. On average, 34.9% of A549 cells displayed a red signal, confirming their mitochondrial-hijacking activity (**Figures 1B and 1C**). Extending this result to all the other cell lines, we observed higher red fluorescent signals in cocultured cancer cells than monocultured control and confirmed that up to 40% of cancer cells hijacked mitochondria from T cells *in vitro* (**Figures S1C-F**). To assess if mitochondrial transfer is mediated via TNTs, we used a Transwell system to block direct cell–cell contact while still permitting the exchange of extracellular vesicles (EVs). Indeed, the transfer was abolished with pore size small enough to prohibit T cell passing (**Figures S2A and S2B**), which confirmed that TNT is the dominant channel for mitochondria transfer between cancer and T cells.

**Figure 1.**
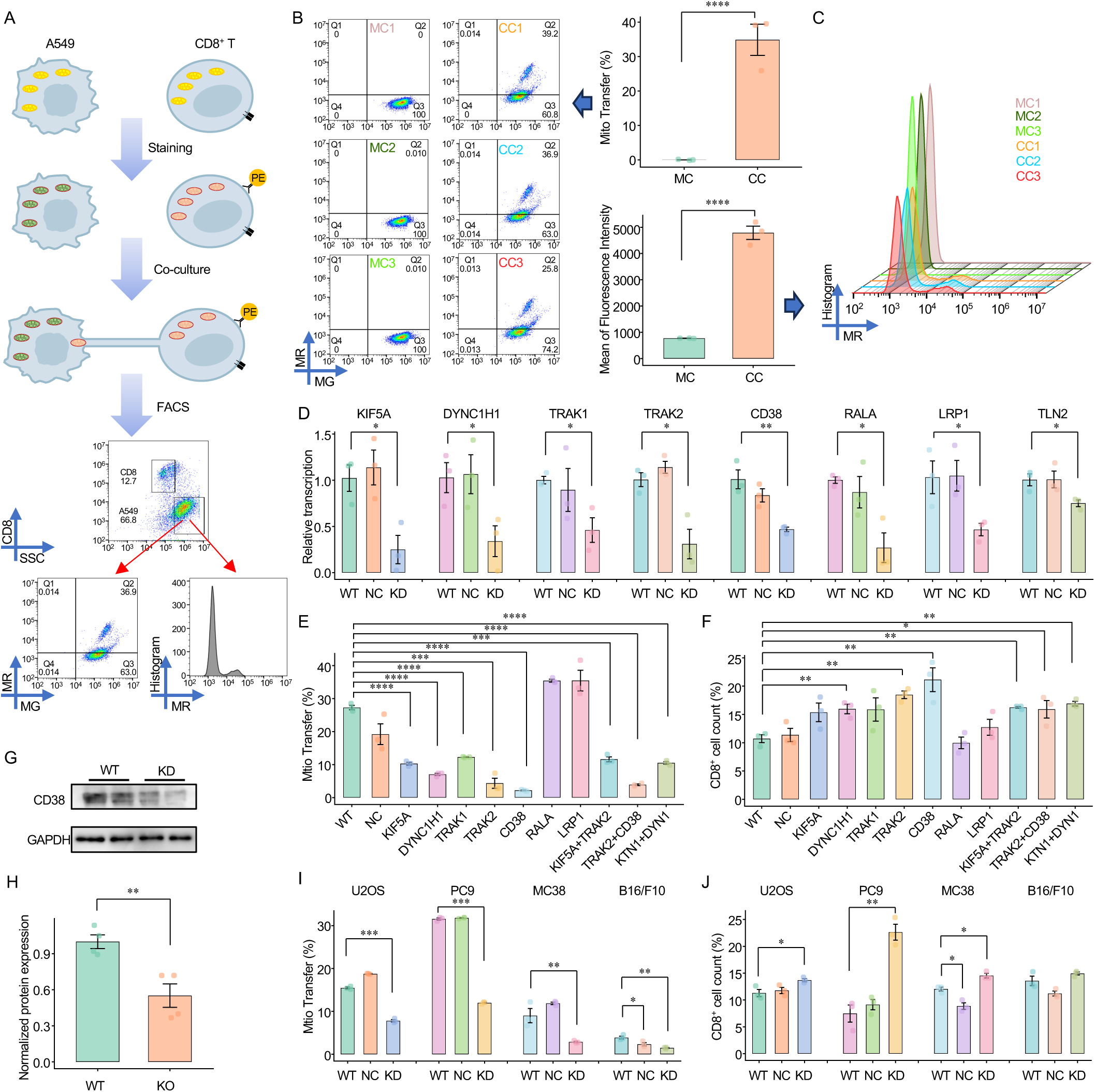
Target screen for blocking intercellular mitochondrial transfer. (A) Schematic overview of the experimental workflow for detecting mitochondrial transfer from T cells to cancer cells. (B) Flow cytometry analysis of the proportion of A549 cancer cells acquiring T cell–derived mitochondria (MitoTracker Deep Red, MR) under mono-culture or co-culture conditions for 3 h. **** *P* < 0.0001. Results are the mean ± SE (n = 3). (C) Flow cytometry analysis of MR fluorescence intensity in A549 cancer cells cultured alone or co-cultured with T cells for 3 h. **** *P* < 0.0001. Results are the mean ± SE (n = 3). (D) mRNA expression levels of motor protein genes (KTN1, DYN1, TRAK1, TRAK2) and TNT-associated genes (CD38, RALA1, LRP1, TLN2) in wild-type (WT), negative control (NC), and shRNA knockdown (KD) A549 cells. * *P* < 0.05, ** *P* < 0.01. Results are the mean ± SE (n = 3). (E) Proportion of WT and KD A549 cancer cells acquiring T cell–derived mitochondria, measured by flow cytometry. *** *P* < 0.001, **** *P* < 0.0001. Results are the mean ± SE (n = 3). (F) Proportion of CD8 T cells following 3 h co-culture. * *P* < 0.05, ** *P* < 0.01. Results are the mean ± SE (n = 3). (G) Immunoblot analysis of CD38 protein expression in WT and CD38 KD A549 cells. (H) Quantification of CD38 protein levels shown in (G). ** *P* < 0.01. Results are the mean ± SE (n = 4). (I) Proportion of WT, NC, and CD38 KD cancer cells (U2OS, PC9, MC38, and B16/F10) acquiring T cell–derived mitochondria, measured by flow cytometry. * *P* < 0.05, ** *P* < 0.01, *** *P* < 0.001. Results are the mean ± SE (n = 3). (J) Proportion of CD8□ T cells following 3 h co-culture. * *P* < 0.05, ** *P* < 0.01. Results are the mean ± SE (n = 3).

A number of genes/pathways have been reported to influence mitochondria trafficking. To determine which one(s) affect cancer-mediated mitochondrial hijack of T cells, we performed a thorough literature search and selected the top candidates involved in mitochondria trafficking or nanotube regulation: *RALA*, small GTPase that triggers TNT formation^13,14^; *LST1*, upstream regulator of *RALA*^15^; *CD38*, surface glycoprotein facilitates TNT formation^16^; *TLN2*, a cytoskeletal protein that mediates membrane protrusions^17^; *LRP1*, an endocytic/signaling cell-surface receptor that mediates astrocyte-to-neuron mitochondrial transfer via extracellular vesicles (EVs)^18^; *KIF5A, DYNC1H1, TRAK1,* and *TRAK2* are motor proteins that mediate intracellular mitochondrial transport^19^. As some of these genes are essential, we applied RNA interference (RNAi)–mediated knockdown (KD), and successful reduced the transcript level for each target gene with shRNA (**Figure 1D**). Notably, for most of the targets, RNAi-mediated silencing did not significantly increase cell apoptosis (**Figure S2C**).

### *CD38* knockdown effectively blocks mitochondria transfer

We next measured mitochondrial transfer under each KD condition. RNAi reduced mRNA levels of *RALA* and *LRP1* by 70%, yet the transfer between A549 cells and T cells was minimally affected (**Figures 1D and 1E**). In contrast, ∼30% knockdown of *TLN2* significantly suppressed the transfer (**Figures 1D and S2D**). However, this reduction was accompanied with elevated apoptosis of A549 cells, suggesting an essential role of *TLN2* in cell survival (**Figure S2E**), and that its inhibition of mitochondrial transfer might be confounded by cell death (**Figures S2F and S2G**).

Knockdown of the motor/adaptor proteins for mitochondria trafficking, including kinesin 1 (*KIF5A*), dynein heavy chain (*DYNC1H1*) and kinesin adaptor proteins (*TRAK1/2*), significantly reduced transfer (**Figure 1E**). Interestingly, of all the targets, *CD38* KD reached the highest suppression rate, causing over 90% reduction (**Figure 1E**). Surprisingly, this performance is even better than the double KD of *KIF5A* and *DYNC1H1*, or *KIF5A* and *TRAK2* (**Figures 1E, S2H, and S2I**). Moreover, simultaneous KD of *TRAK2* and *CD38* did not further increase the inhibitory effect compared with *CD38* KD alone (**Figures 1E and S2H**), suggesting that motor/adaptor protein disruption is dispensable when *CD38* expression is reduced. As a positive control, downregulation of TNT formation using cytarabine (AraC) at 1 µM for 24 h^15^ significantly suppressed mitochondrial transfer (**Figure S2I**). This treatment does not influence cell survival (**Figure S2J**), but is less effective compared to *CD38* KD. These results collectively indicate that *CD38* KD is a sufficient condition to block T cell mitochondria transfer to A549 cells. We confirmed the efficacy of *CD38* KD in the other four cancer cell lines with both human and murine origins (**Figures 1G-J**). This observation may apply to other cancer types as well given the widespread expression of *CD38* in human cancers (**Figure S2K**)^20^.

### *CD38* regulates TNT formation through ligand-binding

To test if *CD38* KD inhibits cancer cell acquisition of T-cell mitochondria by suppressing TNT formation, we co-cultured the WT and *CD38* KD A549 cells with CD8+ primary T cells and stained F-actin within nanotubular structures using phalloidin. Confocal imaging confirmed that *CD38* KD significantly reduced the number and length of TNTs (**Figures 2A-C**).

**Figure 2.**
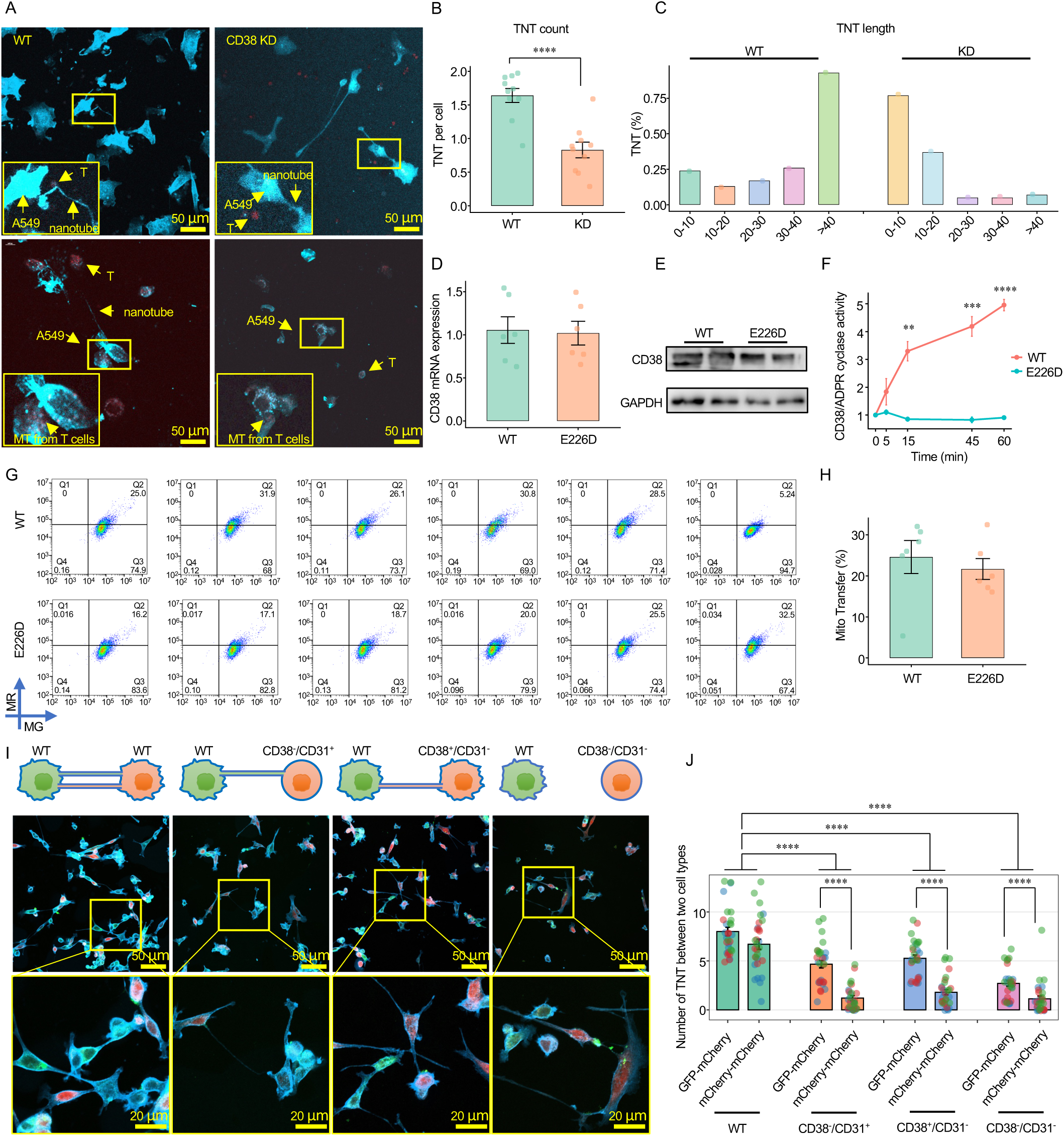
CD38 knockdown inhibits tunneling nanotube (TNT) formation. (A) Confocal imaging of TNTs between A549 cancer cells and T cells under co-culture conditions in wild-type (WT) and CD38 knockdown (KD) groups. Scale bars, 50 μm. (B) Quantification of TNT number per A549 cell in (A). Ten randomly selected fields of view were analyzed, and TNT numbers were calculated on a per-cell basis within each field. Each data point represents one field of view. **** *P* < 0.0001. Results are the mean ± SE (10 fields). (C) Distribution of TNT lengths measured from confocal images under WT and CD38 KD conditions. All identifiable TNT structures were measured and grouped into length intervals (0–10, 10–20, 20–30, 30–40, and >40 μm), and the percentage of TNTs within each interval was calculated. (D) mRNA expression of CD38 in A549 cells following CRISPR-mediated CD38 knockout and reconstitution with WT or E226D mutant CD38. Results are the mean ± SE (n = 6). (E) Immunoblot analysis of CD38 protein expression in A549 cells expressing WT or E226D mutant CD38. (F) ADP-ribosyl cyclase activity in A549 cells expressing WT or E226D mutant CD38. ** *P* < 0.01, *** *P* < 0.001, **** *P* < 0.0001. Results are the mean ± SE (n = 3). (G) Flow cytometry analysis of the proportion of WT and E226D A549 cancer cells acquiring T cell–derived mitochondria following 3 h co-culture. (H) Quantification of mitochondrial transfer shown in (G). Results are the mean ± SE (n = 6). (I) Confocal imaging of TNTs between WT, CD38 KD and CD31 KD A549 cells. Scale bars, 50 μm and 20 μm. (J) Quantification of TNT number in (I). Each color represents an independent biological replicate (n = 3), with 10 randomly selected fields analyzed per replicate (each dot represents one field). **** *P* < 0.0001. Results are the mean ± SE.

Studies in astrocytes and multiple myeloma have largely attributed CD38-dependent mitochondrial transfer to CD38–cADPR–calcium signaling^16,21^. To test if this mechanism is important for T cell mitochondrial loss, we re-expressed a mutant form of the protein, CD38^E226D^ in the CD38-knockout A549 cells. Replacement of Glutamic Acid at 226^th^ position by Aspartic Acid abolishes the enzymatic activity of *CD38*^22^. Consistently, we observed a robust suppression of ADPR cyclase activity without affecting *CD38* mRNA or protein expression levels (**Figures 2D-F**). Interestingly, this enzymatical inactivation of *CD38* had no effect on the ability of A549 cells to acquire mitochondria from T cells (**Figures 2G and 2H**). We further validated this finding in murine MC38 cells by introducing an analogous point mutation at residue 230 of *Cd38* in CD38-KD cells, which similarly failed to reduce mitochondrial transfer (**Figures S3A-C**). Together, these results indicate that, unlike previous observations made for other cell types, CD38-mediated mitochondrial transfer between cancer and T cells is independent of its enzymatic activity.

*CD38* also functions as a ligand-binding molecule, most prominently through its interaction with *CD31* (*PECAM1*), a putative endothelial marker^23^. Based on GTEx data, *CD38* is highly expressed in lymphocytes and lung epithelial cells (**Figures S3D and S3F**) with *CD31* expressed in lung and breast tissues (**Figures S3E and S3G**). To explore whether the *CD38/CD31* axes quantitatively influences TNT connectivity, we expressed fluorophores in the A549 cells with two colors: A549^GFP^ (WT) and A549^mCherry^ (perturbed). Plasma membrane was stained with phalloidin to visualize the TNTs. We knocked down *CD38* and/or *CD31* in A549^mCherry^ cells and cocultured them with WT to measure TNT formation in confocal images (**Figures 2I and 2J**). Specifically, TNTs formed between WT and perturbed or within perturbed cells were calculated. Single KD of *CD38* or *CD31* lead to similar level of reduction of TNT count, whereas double KD further decreased TNT formation between WT and KD cells. In contrast, TNT formation between KD cells is rarely observed (**Figure 2J)**. These results were further quantified using flow cytometry to measure mitochondrial transfer under the same conditions (**Figure S3H**). Together, this semi-quantitative assay supported that *CD38* regulates TNT formation through binding to its cognate ligand *CD31*.

### CD8^+^ T cell dependent suppression of tumor growth by *CD38* silencing

We next investigated how TNT blockade influenced tumor growth *in vivo*. 10^6^ WT (vehicle) cells or *Cd38* knocked down MC38 cells were subcutaneously xenografted to the syngeneic C57BL/6 mice (**Figures S4A and S4B**). Unexpectedly, MC38 cells with *Cd38* KD exhibited markedly impaired tumor growth, reaching only ∼300 mm³ by day 23 (**Figures S4C-E**), suggesting an important role of TNT connectivity during the tumor seeding phase. To allow tumor growth, we constructed a Tet-On shRNA system for conditional *Cd38* KD, where shRNA expression is activated with doxycycline (Dox) (**Figure S4F**). We verified *Cd38* KD by treating MC38 cells with Dox *in vitro*, and the results confirmed that the inducible system functioned efficiently (**Figure S4G**).

Using this system, we grafted 10^6^ WT or Cd38^Tet-On^ MC38 cells to first allow uninterrupted tumor growth. Dox was orally delivered on day 14 for both groups and significant tumor reduction was observed starting from day 29 (**Figures 3A-C**). *Cd38* expression in the tumors collected at day 35 was significantly reduced in the treatment group (**Figure S4H**). The total mitochondrial content in the WT tumors (CD45^-^) is significantly higher than in the *Cd38* KD tumors, confirming the role of mitochondrial transfer blockade by *Cd38* KD *in vivo* (**Figure 3D**). *In vitro* analysis showed that *Cd38* KD did not affect the growth of A549 or MC38 cells (**Figures S4I and S4J**), indicating that the enzymatic function of *CD38* does not influence cancer cell proliferation. To explore if the control of tumor growth is related to T cells, we added another group by treating *Cd38*^Tet-On^ MC38 tumors with anti-CD8 mAb on day −2, 2, 8, 12 and 16 of xenograft to deplete CD8^+^ T cells. We confirmed that CD8^+^ T cell depletion partially abolished the treatment effect of *Cd38* KD *in vivo* and significantly increased tumor size (**Figures 3E-G**). Immunohistology also showed that T cells were restricted in the peripheral region of WT tumors, whereas *Cd38* KD boosted T cell infiltration in to the central zone (**Figures 3H and 3I**). In conclusion, *Cd38* KD controls tumor growth *in vivo* in a CD8^+^ T cell-dependent manner.

**Figure 3.**
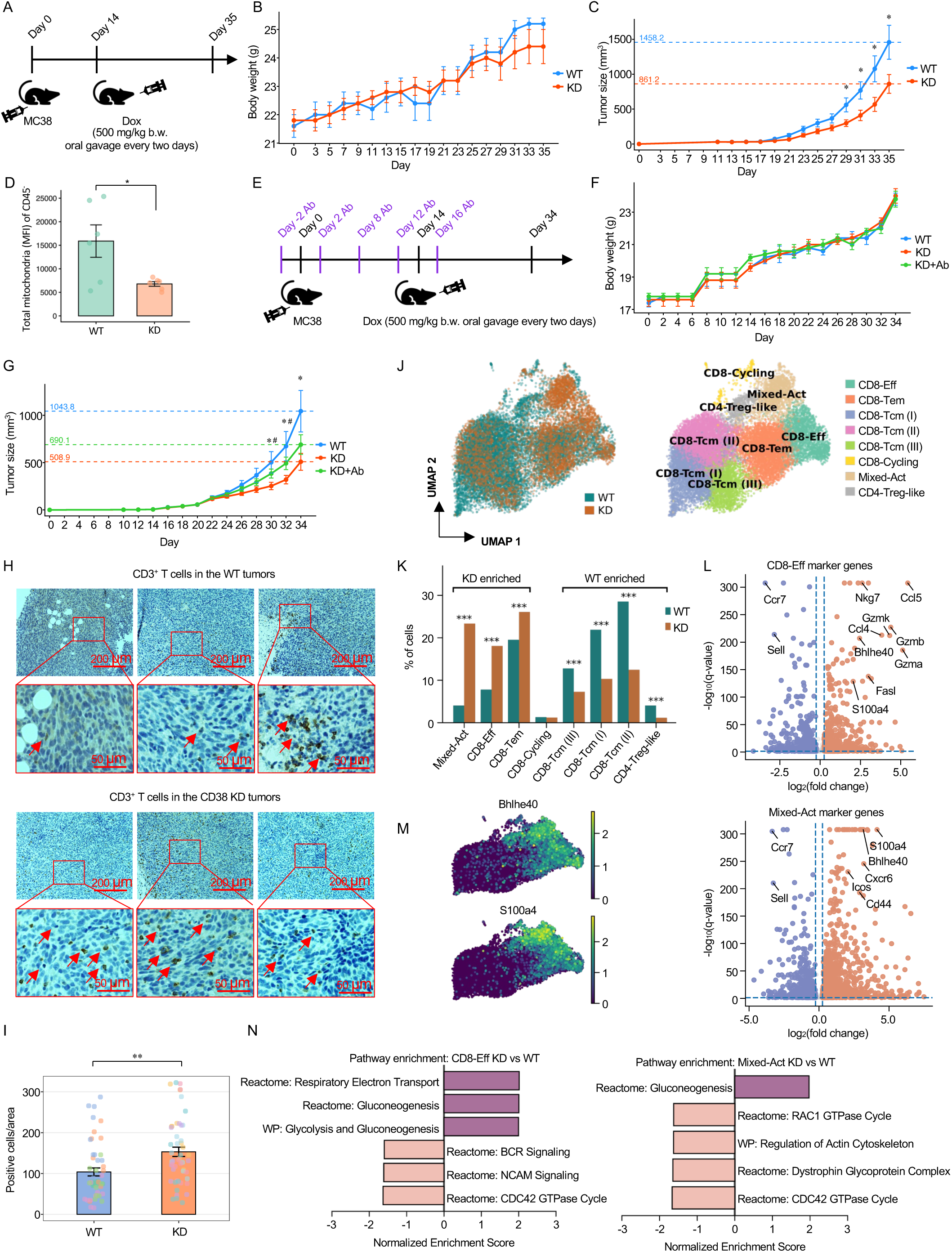
CD8□ T cell–dependent suppression of tumor growth by CD38 silencing. (A) Schematic of the experimental design for subcutaneous implantation of CD38 shRNA Tet-On MC38 tumors. (B) Body weights of tumor-bearing mice. Results are the mean ± SE (n = 5). (C) Tumor growth curves of WT and CD38 KD MC38 tumors. * *P* < 0.05 WT vs. KD. Results are the mean ± SE (n = 9). (D) Mean fluorescence intensity (MFI) of MitoTracker Green in CD45□ tumor cells isolated from tumors in (A), indicating total mitochondrial content. * *P* < 0.05. Results are the mean ± SE (n = 6). (E) Schematic of the experimental design for CD38 shRNA Tet-On MC38 tumor implantation with CD8 T cell depletion. (F) Body weights of tumor-bearing mice with CD8□ T cell depletion. Results are the mean ± SE (n = 5). (G) Tumor growth curves under CD8 T cell depletion. * *P* < 0.05 WT vs. KD, # *P* < 0.05 KD vs. KD+Ab. Results are the mean ± SE (n = 10). (H) Immunohistochemical analysis of CD3□ T cell infiltration in tumor tissues. Each color represents an independent biological replicate (n = 5), with 10 randomly selected fields analyzed per replicate (each dot represents one field). ** *P* < 0.001. Results are the mean ± SE. Scale bars, 200 μm and 50 μm. (I) Quantification of CD3^+^ T cell number in (H). Each color represents an independent biological replicate (n = 5), with 10 randomly selected fields analyzed per replicate (each dot represents one field). ** *P* < 0.001. Results are the mean ± SE. (J) UMAP visualization of T cell populations from WT and KD tumors. Five mice per group were analyzed, comprising 12,270 T cells (WT) and 4,911 T cells (KD). (K) Proportions of T cell subsets in WT and KD tumors. Statistical significance was determined by Fisher’s exact test. ***FDR < 0.001. (L) Volcano plots showing differentially expressed genes (DEGs) in CD8-Eff and Mixed-Act clusters compared to other T cell subsets. (M) Log-normalized expression of marker genes in KD-enriched clusters, including Bhlhe40 and its downstream target S100a4. (N) Gene set enrichment analysis (GSEA) of top pathways enriched in CD8-Eff and Mixed-Act clusters in KD versus WT tumors. FDR < 0.05.

### Selective restoration of Bhlhe40+ T cells upon *Cd38* silencing

To characterize T-cell phenotypic changes upon mitochondrial-loss, we performed droplet-based single-cell RNA sequencing of tumor-infiltrating T cells from WT and *Cd38* KD tumors. We obtained a total of 17,181 T cells after quality control and identified eight clusters by pooling all the samples (**Figures 3J, S5A, and S5B**). Comparative analysis revealed significant enrichment of two T cell subsets in the *Cd38* KD group compared to WT tumors: CD8^+^ effector (CD8-Eff) and a population mixed with both activated CD4^+^ and CD8^+^ T cells (Mixed-Act), while Treg-like CD4^+^ T cells were depleted (**Figure 3K**). The CD8-Eff cluster exhibited high expression of cytotoxic effector molecules (*Gzmb*, *Gzmk*, *Gzma*, *Nkg7*, *Fasl*) and chemokines (*Ccl5*, *Ccl4*). The Mixed-Act cluster was characterized by activation-associated markers of helper T cells (*Cd44*, *Icos*, *Cxcr6*; **Figures 3L and S5B**). Notably, both clusters highly expressed Basic helix loop helix family member E40 (*Bhlhe40*) among the top upregulated genes (**Figures 3L and 3M**). *Bhlhe40* is a transcription factor that regulates T-cell activation and effector differentiation, which also maintains T cell mitochondria fitness^24^.

The transcriptional programs of the CD8-Eff and Mixed-Act clusters in WT tumors exhibited evidence of mitochondrial loss, when compared to the *Cd38* KD group. Specifically, KD group presented higher metabolic activity, marked by upregulation of genes involved in mitochondrial oxidative phosphorylation (*Ndufb1*, *Cox7c*, *Atp5d*) and glycolysis (*Gapdh*, *Ldha*, *Aldoa*), accompanied by downregulation of genes associated with actin remodeling and cytoskeletal dynamics (*Dock2*, *Elmo1*, *Itga4*, *Itgb1*, *Actb*) (**Figure S5C**). Consistently, we observed significant enrichment of genes in the oxidative phosphorylation pathways in the KD group (**Figure 3N, Table S1 and S2**). These results suggested that cancer cells selectively hijack mitochondria from Bhlhe40^+^ T cells, leading to an altered metabolic program in these cells.

### *BHLHE40* upregulates *S100A4* to permit T cell mitochondria loss

To investigate the potential mechanism underlying the selective vulnerability of Bhlhe40^+^ T cells, we examined the other top upregulated genes in the CD8-Eff cluster and identified *S100a4* as a co-expressed marker (**Figures 3L and 3M**). *S100a4* expression was highly correlated with *Bhlhe40* expression across single cells (Pearson r=0.47, *p*<10^-300^) (**Figure S5D**). *S100a4* is a secreted chemoattractant that guides the membrane protrusions during TNT formation^25^. Although it is a putative target for cancer metastasis^26^, *S100a4* is also highly expressed in activated T cells^27–29^. However, the regulatory mechanisms for *S100a4* expression in T cells remains unclear. We analyzed public ChIP-seq samples and discovered that *BHLHE40* directly binds the *S100A4* promoter region in multiple human lymphocyte cell lines (**Figure 4A**). We further confirmed that in primary human T cells, overexpression (OE) of *BHLHE40* leads to *S100A4* upregulation (**Figure 4B**). *In vitro* assays revealed that *BHLHE40* OE in primary T cells cocultured with WT A549 cells increased mitochondrial hijack by the cancer cells, which is reduced by KD of T cell *S100A4* (**Figures 4C-E)**. Consistently, *S100A4* KD partially rescued the cancer-mediated mitochondrial loss in *BHLHE40* overexpressed T cells (**Figures 4C-E**). We further confirmed that mitochondrial transfer regulated by *BHLHE40* and *S100A4* is primarily mediated by TNTs rather than EVs using Transwell (**Figure S5E**).

**Figure 4.**
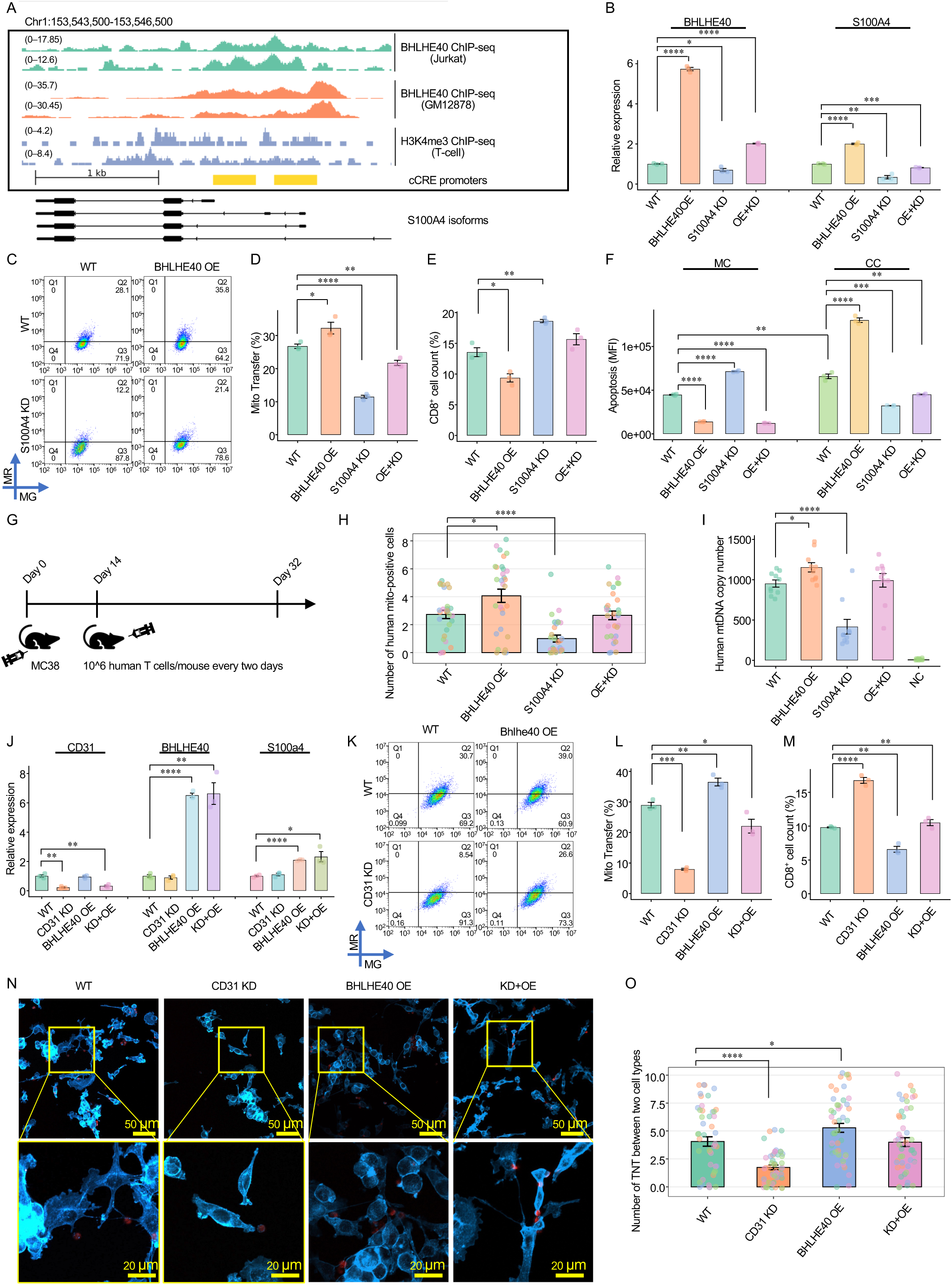
BHLHE40 upregulates S100A4 to promote T cell mitochondria loss. (A) ChIP–seq analysis showing BHLHE40 binding at the promoter region of S100A4. cCRE, ENCODE candidate cis-regulatory elements. (B) mRNA expression of BHLHE40 and S100A4 in T cells under the indicated conditions: wild-type (WT), BHLHE40 overexpression (BHLHE40 OE), S100A4 knockdown (S100A4 KD), and combined BHLHE40 overexpression with S100A4 knockdown (OE+KD) * *P* < 0.05, ** *P* < 0.01, *** *P* < 0.001, **** *P* < 0.0001. Results are the mean ± SE (n = 3). (C) Flow cytometry analysis of mitochondrial transfer to A549 cancer cells co-cultured with WT, BHLHE40 OE, S100A4 KD, or OE+KD T cells. (D) Quantification of mitochondrial transfer shown in (C). * *P* < 0.05, ** *P* < 0.01, **** *P* < 0.0001. Results are the mean ± SE (n = 3). (E) Quantification of CD8□ T cell numbers in (C). * *P* < 0.05, ** *P* < 0.01. Results are the mean ± SE (n = 3). (F) Apoptosis of T cells under the indicated conditions. ** *P* < 0.01, *** *P* < 0.001, **** *P* < 0.0001. Results are the mean ± SE (n = 3). (G) Schematic of the experimental design for subcutaneous implantation of MC38 tumors. A total of 1 × 10□ human T cells (WT, BHLHE40 OE, S100A4 KD, or OE+KD) were injected subcutaneously between two tumors on the dorsal side of mice every two days. (H) Quantification of Hs-MT-No-XMm-positive mouse tumor cells in the tumor sections in Figure S6A. Each color represents an independent biological replicate (n = 6). For each replicate, six randomly selected fields were analyzed, and five human primary T cells were randomly selected from each field for quantification. * *P* < 0.05, **** *P* < 0.0001. Results are the mean ± SE. (I) Human mtDNA levels in mouse CD45□ tumor cells isolated from tumors in (G). NC represents mouse CD45 cells isolated from WT tumors in Figure 3A. * *P* < 0.05, **** *P* < 0.0001. Results are the mean ± SE (n = 10). (J) mRNA expression of CD31, BHLHE40, and S100A4 in T cells under the indicated conditions: WT, CD31 knockdown (CD31 KD), BHLHE40 overexpression (BHLHE40 OE), and combined CD31 knockdown with BHLHE40 overexpression (KD+OE). * *P* < 0.05, ** *P* < 0.01, **** *P* < 0.0001. Results are the mean ± SE (n = 3). (K) Flow cytometry analysis of mitochondrial transfer to A549 cancer cells co-cultured with WT, CD31 KD, BHLHE40 OE, or KD+OE T cells. (L) Quantification of mitochondrial transfer shown in (H). * *P* < 0.05, ** *P* < 0.01, *** *P* < 0.001. Results are the mean ± SE (n = 3). (M) Quantification of CD8□ T cell numbers in (H). * *P* < 0.05, ** *P* < 0.01, **** *P* < 0.0001. Results are the mean ± SE (n = 3). (N) Confocal imaging of TNT formation between A549 cells and T cells under the indicated conditions: WT, CD31 KD, BHLHE40 OE, and KD+OE. Scale bars, 50 μm and 20 μm. (O) Quantification of TNT number in (N). Each color represents an independent biological replicate (n = 3), with 10 randomly selected fields analyzed per replicate (each dot represents one field). * *P* < 0.05, **** *P* < 0.0001. Results are the mean ± SE.

Compared to monocultured primary T cells, coculture with WT cancer cells caused elevated T cell death (**Figure 4F**). Paradoxically, *BHLHE40* OE under monoculture condition improved T cell survival, yet significantly increased T cell apoptosis when cocultured with WT A549 cells. Likewise, while *S100A4* inhibition induced T cell apoptosis when cultured alone, it significantly improved T cell survival under coculture (**Figure 4F**). Metabolic assays revealed that when cocultured with A549 cells, though total ATP production was unchanged, T cells with *BHLHE40* OE exhibited significant increase in Reactive Oxygen Species (ROS) production **(Figures S5F-H)**. Based on these observations, we concluded that the *BHLHE40 → S100A4* cascade promotes cancer-mediated mitochondrial loss in T cells *in vitro*.

### *In vivo* evidence that BHLHE40 drives mitochondrial transfer

We next designed experiments to test if this mechanism occurs *in vivo*. As mitochondrial transfer is cross-species conserved^30^, mouse tumors can acquire mitochondria from human T cells, thus allowing *in vivo* tracing of human mitochondrial transfer in mouse tumors. To achieve this goal, we synthesized fluorescence in situ hybridization (FISH) probes for human *CD3E* to identify T cells, and a human-specific mitochondrial probe (Hs-MT-No-XMm), to detect human mtDNA without cross-reactivity to mouse mitochondrial DNA. We then injected primary human T cells with 1) no treatment (WT), 2) *BHLHE40* OE, 3) *S100A4* KD and 4) *BHLHE40* OE + *S100A4* KD (OE + KD) next to the subcutaneously grafted MC38 tumors (**Figure 4G**). Consistent with our expectation, Hs-MT-No-XMm^+^ cells were detected near T cells in the tumor regions, confirming *in vivo* T cell to cancer mitochondrial transfer (**Figure S6A**). Quantitative assay further confirmed that *BHLHE40* OE markedly increased the number of neighboring mouse tumor cells positive for human mtDNA, whereas *S100A4* KD reduced the transfer to close T cell vicinity despite comparable infiltration of human T cells into the tumors **(Figure 4H)**.

As FISH only accesses thin tissue slices, we sought to confirm the above findings by using a human mtDNA-specific qPCR assay to quantify mtDNA transfer of the whole tumor. This assay leveraged an independent set of primers to selectively amplify the human mitochondrial genome while excluding the nuclear mitochondrial DNA elements (NUMTs). We confirmed the primer specificity using human A549 cells and mouse MC38 cells **(Figure S6B)**, and isolated CD45 cells from mouse tumors to exclude infiltrating human T cells. In line with our previous observations, human mtDNA was readily detected in mouse tumor cells, with *BHLHE40* OE significantly increasing mtDNA transfer. Consistently, *S100A4* KD reduced transfer by over 60% **(Figure 4I)**. Together, these findings demonstrated that *BHLHE40 → S100A4* cascade promotes mitochondrial transfer from T cells to tumor cells *in vivo*. Notably, in the OE+KD group, *S100A4* KD cannot completely offset mitochondrial loss by *BHLHE40* overexpression (**Figure 4H-I**).

This is likely because in OE+KD T cells, *BHLHE40* OE compensated *S100A4* expression to nearly WT level (**Figure 4B**), leading to similar extent of mitochondrial loss as in the WT group. Our results also indicated that human *S100A4* retains its biological activity in the murine tumor microenvironment, which is likely due to the high sequence conservation between human *S100A4* and mouse *S100a4* **(Figure S6C)**.

### *BHLHE40* OE compensates *CD31* downregulation for mitochondrial transfer in T cells

We next investigated how *CD31* and *BHLHE40* OE jointly influence mitochondria transfer. *CD31*, or *PECAM1* is highly expressed in naïve T cells, yet reduced after T cell activation^31^. Although low *CD31* expression was observed in *ex vivo* activated T cells (**Figure S7A**), this downregulation did not prevent mitochondrial transfer (**Figures S7B-D**). In contrast, activated T cells remain vulnerable to cancer-mediated hijacking, likely due to the upregulation of *BHLHE40* and *S100A4* (**Figures S7E and S7F**). To test this hypothesis, we generated CD31-knockdown human T cells. Notably, *CD31* KD had no effect on *S100A4* expression, and *BHLHE40* OE did not alter *CD31* level (**Figure 4J**). Coculture of *CD31* KD T cells with A549 cells revealed significantly reduced mitochondrial hijack by cancer cells (**Figures 4K-M**), whereas *BHLHE40* OE partially restores mitochondrial transfer despite *CD31* KD (**Figures 4K-M**).

To delineate how *BHLHE40* OE functionally overrides the effects of *CD31* KD, we performed high-resolution confocal imaging of A549–T cell cocultures. *CD31* KD markedly reduced the ability of T cells to form TNT connections, with a considerable fraction of T cells remaining completely disconnected from cancer cells. In contrast, *BHLHE40* OE enhanced the directional extension of TNTs towards T cells, frequently resulting in the connection of multiple TNTs to the same T cell (**Figures 4N and 4O**). In other words, even in the context of *CD31* KD, *BHLHE40* OE partially restored TNT connectivity, enabling multiple TNTs to engage a subset of T cells, while others remained entirely TNT-negative. In summary, we demonstrated how *BHLHE40* overexpression promotes mitochondrial loss in activated T cells with reduced *CD31* expression.

### Neoantigen-specific T cells endure higher mitochondrial-loss

We expect that tumor neoantigen-specific T cells to be more vulnerable to cancer-mediated mitochondrial hijack as they become closely attached to the cancer cells upon TCR/antigen binding^32^. To investigate their behaviors *in vivo*, we designed a tumor-specific lipid nanoparticle (LNP) vaccine targeting the *Adpgk* neoantigen in the MC38 cells^33,34^. The vaccine was administered to mice intravenously on days 10 and 17 post xenograft for both WT and *Cd38* KD tumors **(Figures 5A-C)**. Tumor-infiltrating T cells stained with anti-CD3 antibody and *Adpgk* MHC-I tetramer were isolated by flow cytometry **(Figure S8A)**. Expectedly, LNP vaccination increased the number of the infiltrating Tetramer T cells **(Figure 5D)**, which exhibited stronger apoptotic signals compared to non-specific T cells **(Figure S8B)**. Neoantigen-specific T cells exhibited higher ROS levels **(Figure S8C)**, which is likely caused by elevated mitochondrial-loss. Compared to the *Cd38* KD group, the WT group exhibited a significant reduction in Tetramer T cells, with no difference for the Tetramer group **(Figure 5D)**. We further confirmed that Tetramer T cells expressed significantly higher levels of *Bhlhe40* (**Figure 5E**) and *S100a4* **(Figure 5F)**, with reduced *Cd31* expression (**Figure 5G**).

**Figure 5.**
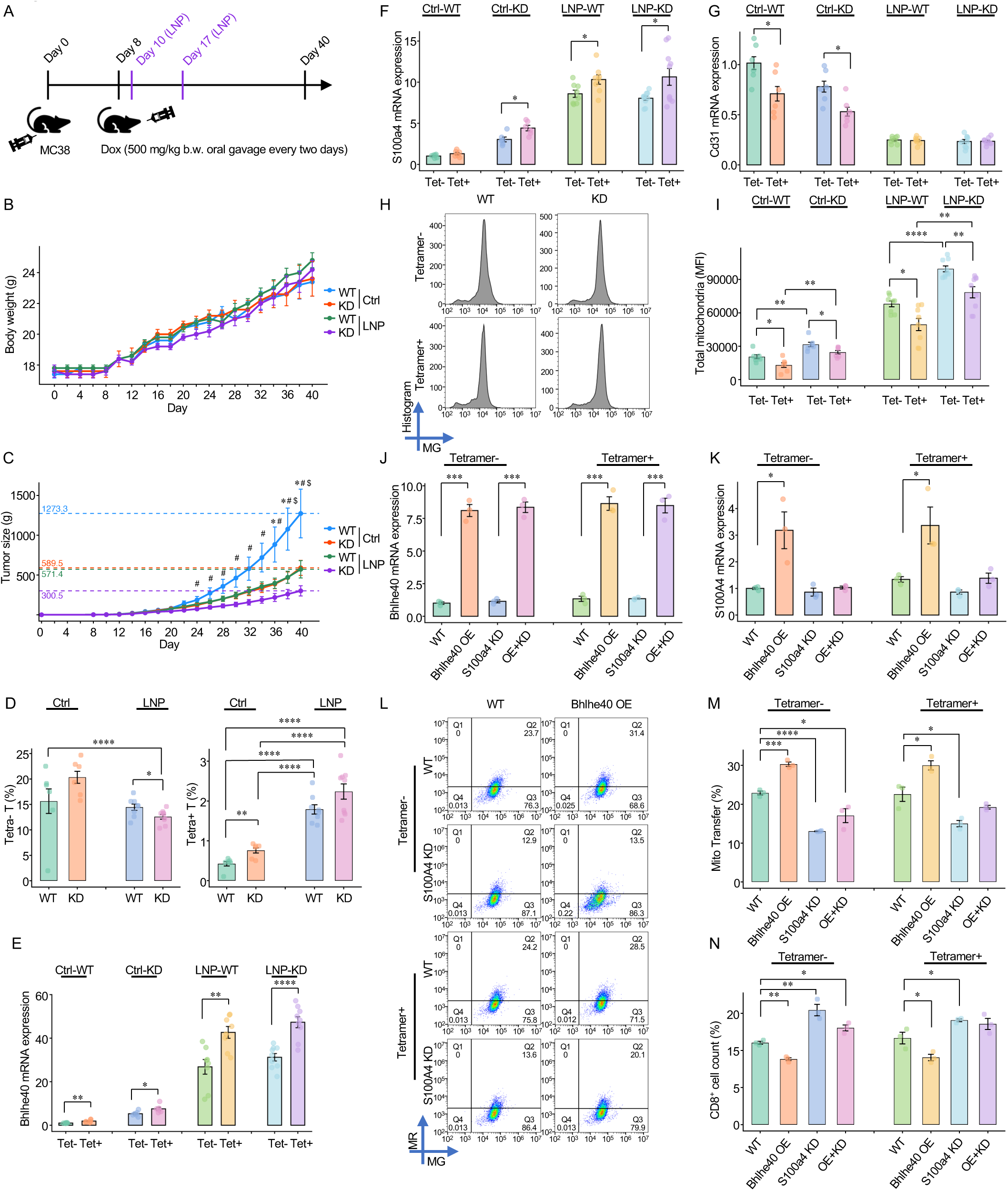
Elevated Bhlhe40 drives T cell mitochondrial loss *in vivo*. (A) Schematic of the experimental design for subcutaneous implantation of CD38 shRNA Tet-On MC38 tumors with lipid nanoparticle (LNP) vaccine treatment. (B) Body weights of tumor-bearing mice. Results are the mean ± SE (n = 10). (C) Tumor growth curves under the indicated conditions. * *P* < 0.05 Ctrl-WT vs. Ctrl-KD, # *P* < 0.05 LNP-WT vs. LNP-KD, $ *P* < 0.05 Ctrl-WT vs. LNP-WT. Results are the mean ± SE (n = 9 tumors for Ctrl-WT, Ctrl-KD, and LNP-WT groups; n = 10 tumors for LNP-KD group). (D) Numbers of tetramer□ and tetramer□ T cells isolated from tumors in (A). * *P* < 0.05, **** *P* < 0.0001. Results are the mean ± SE (n = 6 tumors for Ctrl-WT and Ctrl-KD; n = 8 tumors for LNP-WT; n = 9 tumors for LNP-KD). (E) Apoptosis of tetramer□ and tetramer□ T cells from tumors in (A). ** *P* < 0.01, *** *P* < 0.001, **** *P* < 0.0001. Results are the mean ± SE (n = 6 tumors for Ctrl-WT and Ctrl-KD; n = 8 tumors for LNP-WT; n = 9 tumors for LNP-KD). (F) mRNA expression of Bhlhe40 in tetramer and tetramer T cells isolated from tumors in (A). * *P* < 0.05, ** *P* < 0.01, **** *P* < 0.0001. Results are the mean ± SE (n = 6 tumors for Ctrl-WT and Ctrl-KD; n = 8 tumors for LNP-WT; n = 9 tumors for LNP-KD). (G) mRNA expression of S100a4 in tetramer and tetramer T cells isolated from tumors in (A). * *P* < 0.05. Results are the mean ± SE (n = 6 tumors for Ctrl-WT and Ctrl-KD; n = 8 tumors for LNP-WT; n = 9 tumors for LNP-KD). (H) Flow cytometry analysis of total mitochondrial content in tumor-infiltrating cells isolated from (A). (I) Quantification of mitochondrial content shown in (F). * *P* < 0.05, ** *P* < 0.01, **** *P* < 0.0001. Results are the mean ± SE (n = 6 tumors for Ctrl-WT and Ctrl-KD groups; n = 8 tumors for LNP-WT; n = 9 tumors for LNP-KD). (J) mRNA expression of Bhlhe40 in tetramer□ and tetramer□ T cells isolated from tumors in (A), followed by Bhlhe40 overexpression and/or S100a4 knockdown. *** *P* < 0.001. Results are the mean ± SE (n = 3). (K) mRNA expression of S100a4 under the indicated conditions as in (H). * *P* < 0.05. Results are the mean ± SE (n = 3). (L) Functional assay of tetramer□ and tetramer□ T cells isolated from tumors in (A), followed by Bhlhe40 overexpression and/or S100a4 knockdown, and subsequent co-culture with MC38 cells. (M) Quantification of mitochondrial transfer shown in (J). * *P* < 0.05, *** *P* < 0.001, **** *P* < 0.0001. Results are the mean ± SE (n = 3). (N) Quantification of CD8□ T cell numbers in (J). * *P* < 0.05, ** *P* < 0.01. Results are the mean ± SE (n = 3).

We then tested if neoantigen-specific T cells have higher mitochondrial loss *in vivo* through measuring the total mitochondrial content using MitoTracker fluorescence intensity in the tumor-infiltrating T cells (**Figure 5H**). This assay confirmed that Tetramer T cells contained fewer mitochondria than Tetramer T cells, with *Cd38* KD group exhibiting higher mitochondrial content than WT control **(Figure 5I)**. These results are in line with our previous observation that mitochondrial levels of CD45^-^ tumor cells were significantly higher in the WT group than in the KD group **(Figure 3D)**. Together, our findings confirmed that neoantigen-specific T cells experienced elevated mitochondrial loss in the tumor microenvironment.

### Elevated *Bhlhe40*, instead of antigen-specificity, drives T cell mitochondrial loss *in vivo*

We then asked which factor, T cell antigen specificity, or *Bhlhe40* overexpression, is the cause of higher mitochondrial hijack among the Tetramer+ T cells. To address this, we first collected Tetramer and Tetramer T cells from LNP vaccinated mice and cultured them *ex vivo*. We transfected DNA plasmids to overexpress *Bhlhe40* and/or to knockdown *S100a4*. These T cells were then reactivated and cocultured with MC38 cells. As described above, neoantigen-positive T cells directly isolated from mice exhibited higher expression levels of *Bhlhe40* (**Figure 5E**) and *S100a4* (**Figure 5F**). However, after activation, *Bhlhe40* and *S100a4* expression levels were similar between Tetramer^+^ and Tetramer^-^ T cells by RT-PCR **(Figures 5J and 5K)**. This decoupling of neoantigen-specificity and *Bhlhe40* expression is likely due to *ex vivo* reactivation, yet providing an ideal opportunity to dissect the cause of T cell mitochondrial-loss. We measured mitochondrial transfer from T cells to MC38 cells, and observed that, Tetramer T cells without *Bhlhe40* upregulation experienced similar level of mitochondrial-hijacking by MC38 cells as the non-specific T cells **(Figure 5L)**. *Bhlhe40* OE in T cells elevated transfer to MC38, whereas KD of *S100a4* reduced transfer in the Tetramer^+^ and Tetramer^-^ T cells regardless of *Bhlhe40* levels (**Figure 5M**). We further confirmed that in both Tetramer^+^ and Tetramer^-^ T cells, *Bhlhe40* OE reduced cell viability when cocultured with cancer cells (**Figure 5N**). Together, these results demonstrated that the cascade of *Bhlhe40* overexpression and *S100a4* secretion, instead of tumor neoantigen specificity, plays a dominant role in cancer mediated T cell mitochondria loss.

### Dual mitochondria transfer blockade as an anti-cancer therapy

The above results indicated two potential strategies to prevent cancer cells from hijacking mitochondria in the T cells: 1) block TNT formation via *CD38* knockdown and 2) disruption of TNT guidance by *S100A4* inhibition. We tested both strategies, separately and combined. First, we used a human anti-CD38 monoclonal antibody (Isatuximab) to inhibit *CD38* expression in A549 cells. This treatment (10ug/ml, 24h) markedly suppressed mitochondrial hijacking by cancer cells and protected T cells from damage *in vitro* **(Figures 6A-C)**. Next, we performed *in vivo* experiments in which mice were administered with intraperitoneal injection (i.p.) of trifluoperazine (TFP) daily. TFP is a small molecule inhibitor that disrupts the S100A4–myosin-IIA interaction and reduces the fraction of S100A4 pentamers^35^. We confirmed that TFP significantly suppressed tumor growth (**Figures 6D and 6E**). Finally, we tested the therapeutic effect of dual mitochondrial transfer blockade, with (TFP + *Cd38* KD), and observed more pronounced suppression of tumor growth compared to either treatment alone, indicating a synergistic effect **(Figure 6E)**.

**Figure 6.**
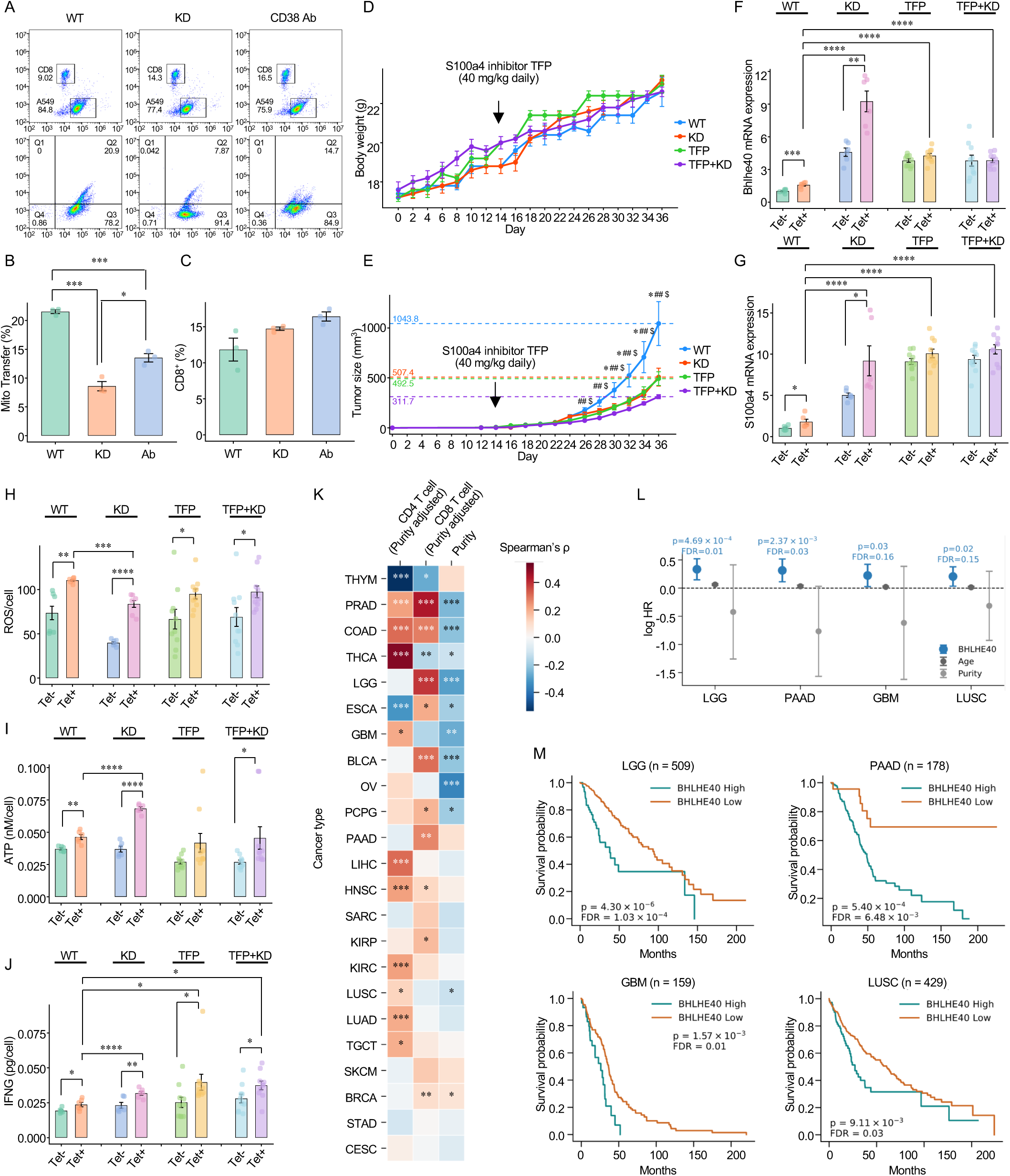
Targeting mitochondrial transfer as a therapeutic strategy for cancer. (A) Effects of Isatuximab antibody treatment (10µg/ml, 24h) on mitochondria transfer in A549 cells. (B) Quantification of mitochondrial transfer shown in (A). * *P* < 0.05, *** *P* < 0.001. Results are the mean ± SE (n = 3). (C) Quantification of T cell numbers in (A). Results are the mean ± SE (n = 3). (D) Body weights of mice bearing CD38 shRNA Tet-On MC38 tumors treated with the S100a4 inhibitor trifluoperazine (TFP) via oral administration. Results are the mean ± SE (n = 10). (E) Tumor growth curves under TFP treatment. * P < 0.05 WT vs. TFP, ## P < 0.01 WT vs. TFP+KD, $ P < 0.05 TFP vs. TFP+KD. Results are the mean ± SE (n = 10). (F) mRNA expression of Bhlhe40 in tetramer and tetramer T cells isolated from tumors in (E). *** *P* < 0.001. Results are the mean ± SE (n = 6 tumors for WT and D; n = 10 tumors for TFP; n = 9 tumors for TFP+KD). (G) mRNA expression of S100a4 in tetramer and tetramer T cells isolated from tumors in (E). * *P* < 0.05. Results are the mean ± SE (n = 6 tumors for WT and D; n = 10 tumors for TFP; n = 9 tumors for TFP+KD). (H) Reactive oxygen species (ROS) levels in tetramer and tetramer T cells isolated from tumors in (E). * *P* < 0.05, ** *P* < 0.01. Results are the mean ± SE (n = 6 tumors for WT and D; n = 10 tumors for TFP; n = 9 tumors for TFP+KD). (I) ATP levels in tetramer□ and tetramer□ T cells isolated from tumors in (E). ** *P* < 0.001, *** *P* < 0.001. Results are the mean ± SE (n = 6 tumors for WT and D; n = 10 tumors for TFP; n = 9 tumors for TFP+KD). (J) IFN-γ (IFNG) production in tetramer□ and tetramer□ T cells isolated from tumors in (E). * *P* < 0.05. Results are the mean ± SE (n = 6 tumors for WT and D; n = 10 tumors for TFP; n = 9 tumors for TFP+KD). (K) Spearman’s correlation between BHLHE40 expression and CD4 T cell fraction, CD8 T cell fraction, and purity across cancer types. * FDR < 0.05, ** FDR < 0.01, *** FDR < 0.001. (L) Hazard ratios (HRs) for BHLHE40 expression and covariates (age or purity) in Cox proportional hazards models. The P-value and FDR for BHLHE40 is shown. BHLHE40 was binarized using the optimal cutpoint procedure, which selects the threshold that maximizes the log-rank statistic. (M) Kaplan-Meier estimates of overall survival. The patients were stratified using the same cutpoints as (L). LGG, Brain Lower Grade Glioma. PAAD, Pancreatic adenocarcinoma. GBM, Glioblastoma multiforme. LUSC, Lung squamous cell carcinoma.

We next assessed the phenotypic changes of Tetramer and Tetramer T cells from tumors under different treatments. First, we confirmed that Tetramer^+^ T cells in *Cd38* KD group expressed significantly higher *Bhlhe40* (**Figure 6F**). Interestingly, mitochondrial transfer blockade therapies generally increased *S100a4* expression in the T cells **(Figure 6G)**. This is consistent with our scRNA-seq data analysis, and is likely caused by higher *Bhlhe40* levels in the tumor-infiltrating T cells (**Figures 3L and 3M**). Functional profiling revealed that *Cd38* KD reduced ROS level and increased ATP production (**Figures 6H and 6I**), while all transfer blockade therapies significantly increased interferon-expression in the neoantigen-specific T cells **(Figure 6J)**. These results collectively indicate that blocking cancer/T cell mitochondrial transfer effectively controls tumor growth by improving the metabolic and cytotoxic functionalities of tumor-infiltrating T cells.

### Clinical relevance of *BHLHE40* in human cancer patients

We extended efforts to investigate *BHLHE40* expression in human tumor samples in the Cancer Genome Atlas (TCGA) cohort. First, we confirmed that in prostate, colon, brain, esophageal, bladder, ovarian and lung cancers, *BHLHE40* expression is negatively correlated with tumor purity estimated from genomic alterations^36^ (**Figure 6K**). This result suggested that *BHLHE40* transcripts were mainly contributed by cells in the tumor microenvironment with euploid genome. We further confirmed that in these cancer types, as well as pancreatic, kidney and head and neck cancers, *BHLHE40* is positively associated with CD4^+^ or CD8^+^ T cell infiltration level after correction for tumor purity, which indicated a T cell source of the gene expression^37^ (**Figure 6K**). Survival analysis in these cancer types revealed that *BHLHE40* expression negatively predicts patient outcomes for pancreatic cancer, lung squamous carcinoma, lower grade glioma and glioblastoma, after correction for age and tumor purity (**Figures 6L and 6M**). These results suggested that, although *BHLHE40* positively regulates T cell fitness, its contribution to T cell mitochondria loss may lead to worse clinical outcomes.

## Discussion

In this study, we elucidated a novel mechanism of tumor immune evasion by linking *BHLHE40*, a key transcription factor of CD8^+^/CD4^+^ effector T cells, to cancer-mediated mitochondrial hijacking. Revisiting the key questions in cancer/T cell mitochondrial transfer, it becomes clear that cancer cells target effector T cells due to the upregulation of the *BHLHE40 → S100A4* cascade to invite tumor nanotube connections. As a chemoattractant, *S100A4* is also expressed by T-regs, yet significantly lower than effector T cells, thus creating a cancer-targeting preference towards the latter. This process occurs regardless of TCR antigen-specificity status, serving as a dominant pathway for T cell mitochondrial-loss vulnerability.

Generally speaking, the mitochondrial transfer process can be classified into contact-independent and contact-dependent pathways. Contact-independent transfer typically involves the release of mitochondria encapsulated within extracellular vesicles (EVs)^18,38^ and contact-dependent transfer requires specialized structures including TNTs^9,39,40^. Consequently, a variety of strategies have been employed to inhibit mitochondrial transfer, including blocking the endocytic/signaling cell-surface receptor *LRP1* to suppress EV–mediated transfer^18^, or inhibiting TNT formation through introducing mutations in the GTPase *RALA*^13,14^, or knocking down *TLN2*, which mediates interactions with integrins and membrane protrusions^17^. However, each of these approaches has inherent limitations. For example, *LRP1* mediated EVs only occur in specific cell types and its KD is dispensable in our mitochondrial transfer assays between lung cancer cells (A549) and T cells. While *TLN2* knockdown effectively inhibited T-cell mitochondrial-loss, this gene is essential for cell survival. Of all the targets examined, we identified *CD38* as a potent mediator of TNT formation which is conserved across multiple human and mouse cancer cell types.

To our best knowledge, there is no study that clearly defined the molecular mechanisms how *CD38* regulated mitochondrial transfer. *CD38* is an ectoenzyme that regulates intracellular Ca²□ signaling through its ADPR cyclase activity. A previous study has shown that astrocytes may release extracellular mitochondria capsules via CD38-mediated Ca²□ signaling, which are subsequently taken up by neurons after stroke; siRNA-mediated suppression of *CD38* reduces mitochondrial transfer, although the underlying mechanism remains unclear^21^. Some reviews thus proposed that, calcium ions are critical for the formation and maintenance of TNTs, as TNTs are composed of F-actin bundles that can be influenced by Ca²□-dependent dynamics control^16,41,42^. However, this theory has not been definitively proven. In our study, in both human and mouse cancer cells, we showed that inhibition of *CD38*’s cADPR catalytic activity had no effect on cancer/T-cell transfer. This is likely due to the highly redundant pathways for calcium regulation. For example, other enzymes, such as *CD157* and *SARM1*, exert similar catalytic functions as *CD38*^43^. Through a semi-quantitative assay, we showed that *CD38* mediates TNT formation primarily through its receptor–ligand interaction with *CD31*^23^. *CD31* KD significantly reduced intercellular TNT connections and mitochondrial transport. Further, while *CD31* is downregulated in activated T cells^31^, the elevated expression of the *BHLHE40 → S100A4* cascade is sufficient to induce mitochondrial-hijacking through formation of more cancer–derived TNTs to the effector T cells.

Previous studies employed pharmacological inhibitors to block cancer cell–mediated hijacking of T-cell mitochondria, thereby suppressing tumor growth^11,16,21^. For example, treatment of cancer cells with L-778,123, a dual farnesyltransferase and geranylgeranyl transferase inhibitor, has been used to control TNT formation^11^, yet its impact over tumor growth was limited. This is likely because disruption of actin polymerization via GTPase inhibition is non-specific for TNTs. On the other hand, our study showed that dual blockade of TNT formation by *Cd38* KD and *S100a4* inhibitor reduced tumor size by over 70%, superior to all the existing approaches. In theory, mitochondrial-transfer blockade synergizes with other immune-boost therapies, such as checkpoint blockade (ICB). We expect future work to combine this approach with ICB, chimeric-antigen receptor (CAR) T cell therapies, or mRNA vaccines to enhance the efficacy of anti-cancer treatments.

There are several limitations of this study. First, while we provided evidence of mitochondrial transfer with or without *Cd38* KD and/or *S100a4* inhibition in animal models, there is no direct proof that this process is mediated by nanotubes *in vivo*. That being said, detection of TNTs in the tumor tissue, while feasible with advanced imaging technologies^11,30^, is conceptually dispensable to our conclusions. Additionally, although *in vitro* assays proved the efficacy by knocking down *S100A4*, its literature-reported role as a chemoattractant was not directed tested in this work. Further, *in vivo* inhibition of *S100A4* using trifluoperazine might incur off-target effects that are not specific to the T cells. Finally, the dual transfer blockade used RNAi knockdown of *Cd38*, as there is currently no murine anti-*Cd38* antibody for this purpose. Luckily, anti-human *CD38* antibody is being tested in clinical trial for multiple myeloma^44^, which can be repurposed to treat other cancer types with confirmed T cell mitochondrial hijacking activity.

In the future, there are several exciting directions that warrant further investigation. First, it will be interesting to study if there is a specific tumor niche where mitochondrial-hijacking preferentially occurs; Second, other surface molecules, besides the *CD38/CD31* axes, may also participate in TNT connection between cancer and T cells, which might require a large-scale genetic screen to identify; Third, it has been reported that cancer cells may exchange mitochondria with other cell types in the microenvironment, such as macrophage, neutrophil and stromal cells^6,45^. More efforts are needed to elucidate the molecular mechanisms governing these processes; Lastly, our study highlights the functional impacts of heterocellular TNT networks for tumor initiation and progression. A systematic survey of the structure, behavior and function of these networks may lead to novel drug targets and therapeutic opportunities.

## Data and code availability

Raw data and metadata of mouse scRNA-seq are available at GEO accession number GSE326615. Raw data of confocal are available at Zenodo: doi.org/10.5281/zenodo.20089033. GTEx data used for the analyses described in this manuscript were obtained from: the GTEx Portal on 09/22/2025 and dbGaP accession number phs000424.v2.p1 on 09/22/2025. Further information and requests for resources and reagents should be directed to the Lead Contact: Dr. Li.

## Acknowledgments

This work is supported by NCI R01 CA258524 (B.L.).

## Author contributions

B.L. designed the study. H.H. performed the experiments. Y.T. analyzed the single-cell RNA-seq data. I.S. and D.W. designed and synthesized the mRNA-LNP vaccine. Z.Y. and T.W. cultured the cells. J.H., H.Z., and X.Y. analyzed data from clinical sample databases. B.L., Y.T., and H.H. wrote the manuscript. All authors read and approved the final manuscript.

## Key resource table

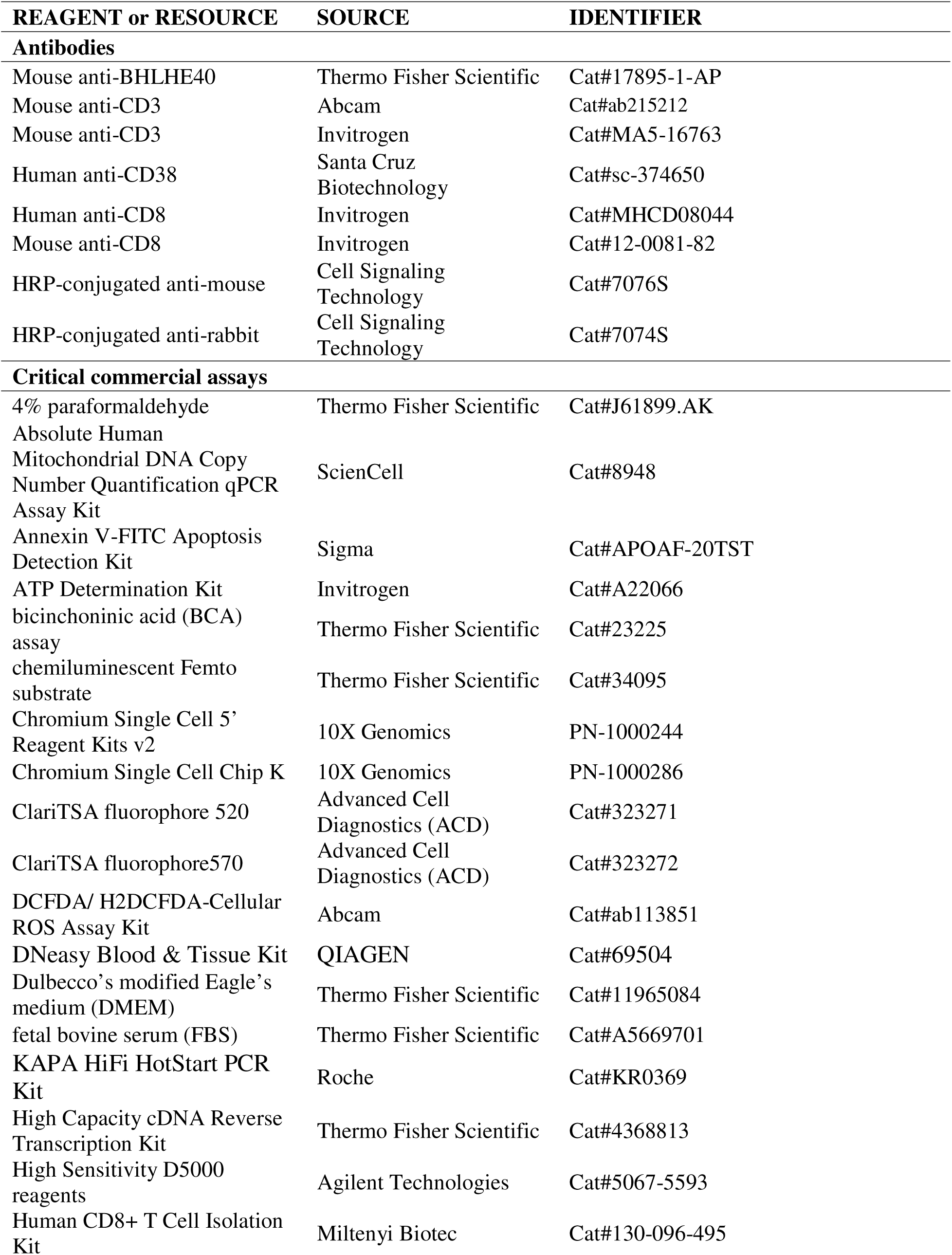
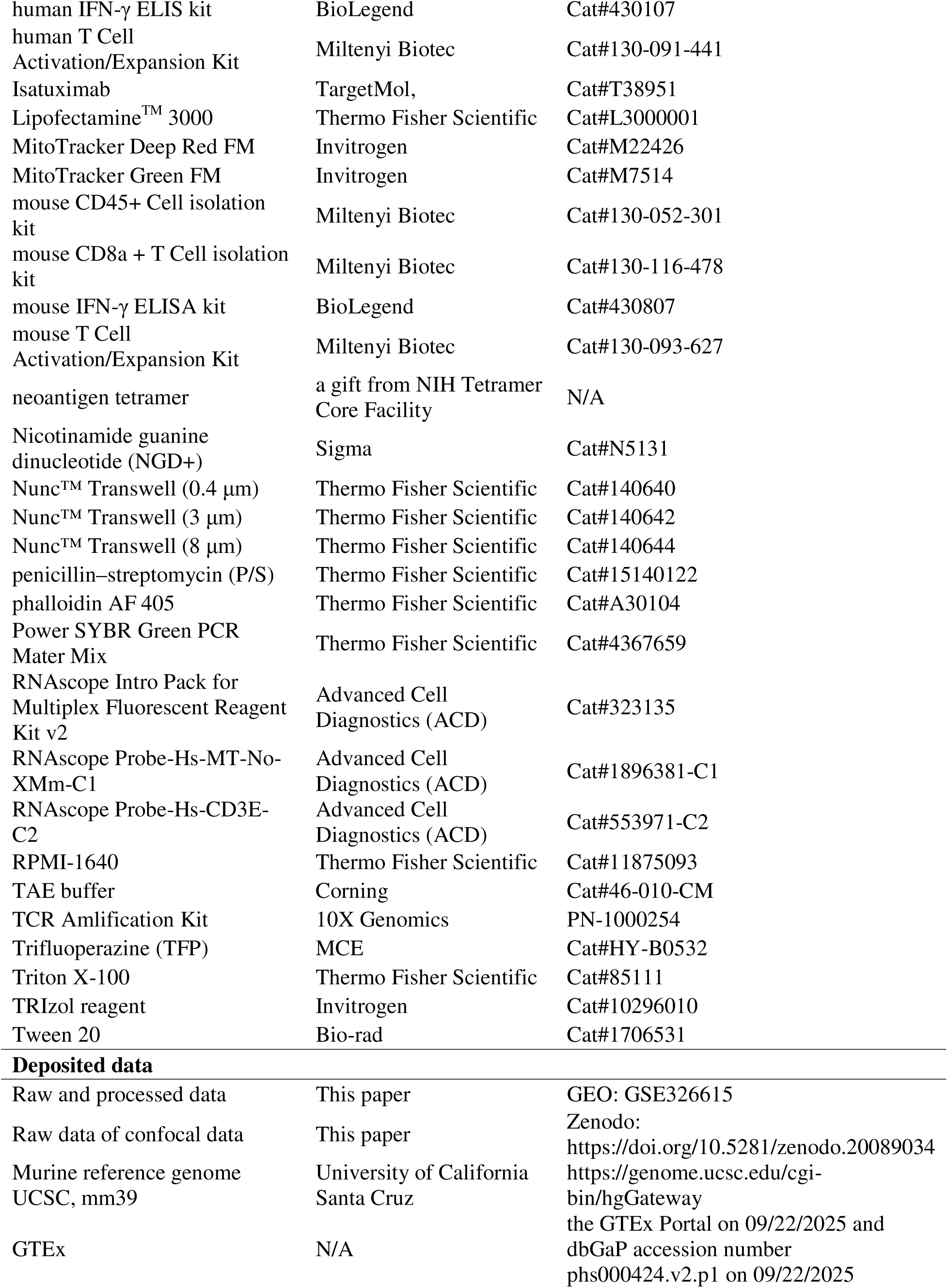
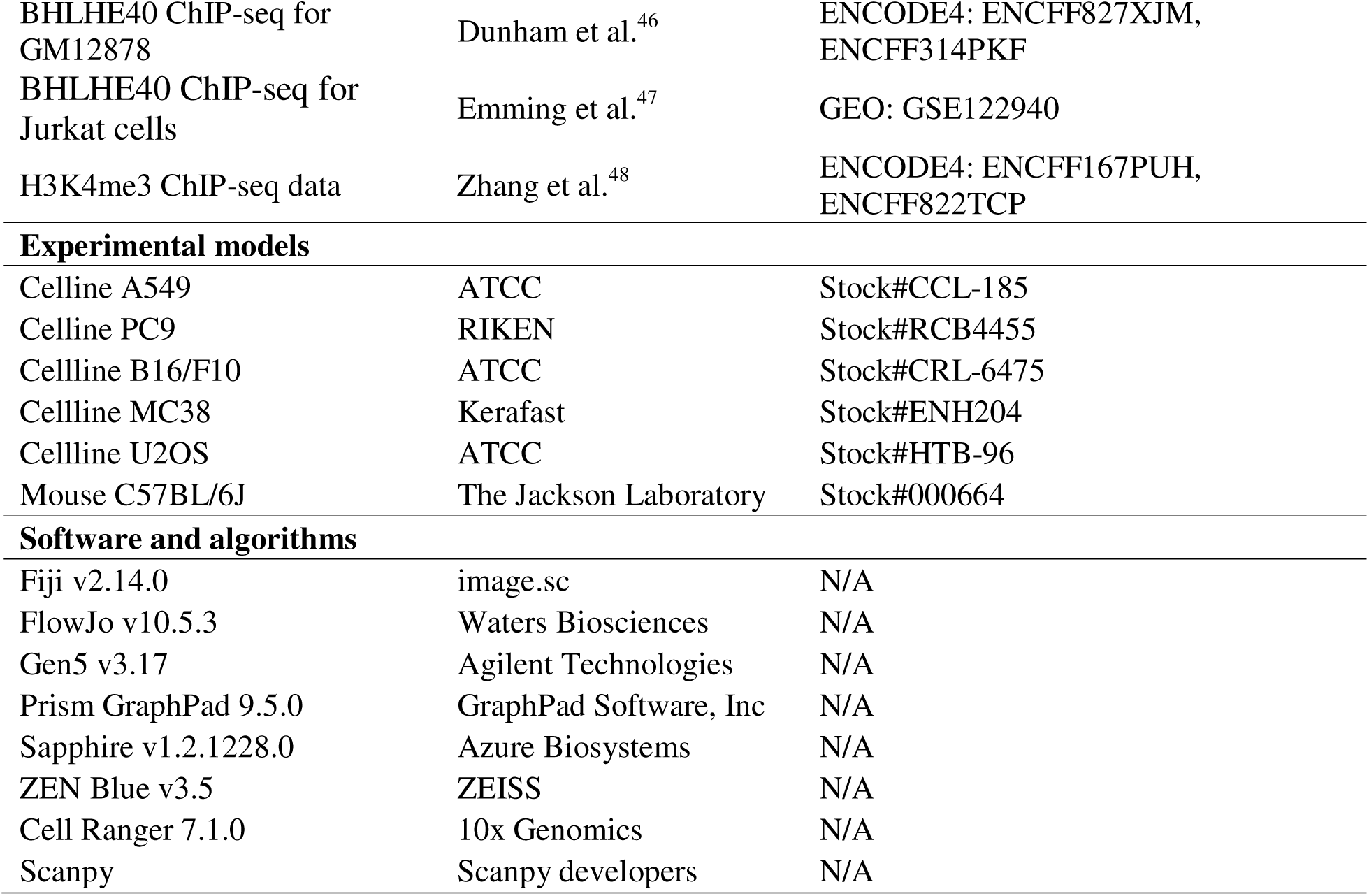

## Materials and Methods

### Cell line culture

A549 (ATCC, #CCL-185); U2OS (ATCC, #HTB-96), PC9 (RIKEN, #RCB4455), MC38 (Kerafast, #ENH204), and B16/F10 (ATCC, #CRL-6475) cells were cultured in Dulbecco’s modified Eagle’s medium (DMEM), supplemented with 10% v/v heat-inactivated fetal bovine serum (FBS) and 1% v/v penicillin–streptomycin (P/S). All cells were cultured at 37 °C in a 5% CO_2_ incubator.

### Human Primary T cells culture

Human primary CD8^+^ T cells were obtained from Human Immunology Core of Perelman School of Medicine, University of Pennsylvania, from healthy volunteers, activated by a human T Cell Activation/Expansion Kit, and cultured in RPMI-1640 supplemented with 10% FBS, 1% HEPES, 1% P/S, under 37 °C in a 5% CO_2_ incubator.

### Gene knockdown and overexpression

Lentiviral and plasmid constructs used in this study included pLKO.1-TRC cloning vector (a gift from David Root; Addgene plasmid #10878), pLKO.1-TCR control (a gift from David Root; Addgene plasmid #10879), Tet-pLKO-puro (a gift from Dmitri Wiederschain; Addgene plasmid #21915), pMD2.G (a gift from Didier Trono; Addgene plasmid #12259), and psPAX2 (a gift from Didier Trono; Addgene plasmid #12260). In addition, pcDNA3-HA-BHLHE40 (a gift from Daniele Guardavaccaro; Addgene plasmid #110154), MSCV-Bhlhe40-3×FLAG-IRES-BFP (a gift from Sarah Teichmann; Addgene plasmid #117263), human wild-type CD38 (VectorBuilder, #VB250910-1368uvc), human CD38 E226D mutant (VectorBuilder, #VB250905-1335zdn), mouse wild-type Cd38 (VectorBuilder, #VB250910-1367qvf), and mouse Cd38 E230D mutant (VectorBuilder, #VB250905-1339uab) were used for gene overexpression studies. Lentiviral particles were produced and amplified in HEK293 cells. Target cells were transduced with lentivirus or LipofectamineTM 3000 for 24–48 hours, followed by selection of transduced cells using puromycin (0.5–10 μg/mL). For inducible knockdown experiments, Tet-pLKO-puro CD38 knockdown cells were treated with doxycycline (1 μg/mL) for 24 hours. The sequences of all shRNAs used in this study are provided in **Table S3**.

### Mice

6-8-week-old female C57BL/6J mice were obtained from Jackson lab and housed under specific pathogen-free conditions at the Children’s Hospital of Philadelphia (CHOP) by following the guidelines of CHOP’s Institutional Animal Care and Use Committee (IACUC). The animals were housed for at least one week before the start of experiments. All the animal procedures were approved by CHOP’s IACUC (protocol #1517). The animals were maintained in dedicated centralized institutional animal facility with a 12 h light/12 h dark cycle. In the rodent housing rooms, the temperature was maintained at 71 °F (±3 °F), and humidity was maintained at 50% (range: 30%–70%).

### Xenograft experiments

Mice were subcutaneously injected in the flanks with wild-type (WT) or CD38 knockdown (KD) MC38 cells (1 × 10^6^ cells per mouse). Each experimental group included five mice, with two tumors established per mouse (one on each flank). Tumor take was not observed in all injection sites; therefore, only tumors that successfully formed were included in subsequent analyses. Final sample sizes are indicated in the corresponding figure legends. Body weight and tumor size were measured every two days. Tumor size was measured by the length (a), width (b) and height (h) and calculated as tumor volume = abh/2. For experiments using Tet-pLKO-puro CD38 plasmid MC38 cells, mice were administered doxycycline (500 mg/kg body weight) by oral gavage every two days starting on Day 8 or Day 14 post-inoculation. For experiments inhibiting S100A4 function *in vivo*, mice were administered 10 mg/kg Trifluoperazine (TFP) daily by intraperitoneal injection (i.p.). Tumors were harvested when they reached a maximum size of 20 mm in any dimension.

### mRNA production and formulation into LNPs

Codon-optimized insert encoding fragment of ADPGK mRNA was cloned into an expression vector for in vitro transcription (IVT) and low endotoxin plasmid DNA Maxiprep was produced by GenScript (Piscataway, NJ). Nucleoside-modified mRNA was produced from linearized DNA templateusing the MegaScript T7 Transcription kit. Uridine was completely substituted for N1-methylpseudouridine (TriLink) and co-transcriptional capping was performed using Clean Cap reagent AG (3’Ome) (TriLink). mRNA was transcribed to include 101 nucleotide long poly(A) tail. After synthesis mRNA was purified of double-stranded mRNA products using cellulose purification^49^. The mRNA sequence is provided in **Table S4**. The ADPGK mRNA expression was confirmed by transfection into HEK293 and B16 cells using Lipofectamine 3000, followed by qPCR.

Purified mRNAs were encapsulated into four-component LNPs using a self-assembley process of ethanolic lipid mixture of an ionizable cationic lipid with 1,2-distearoyl-sn-glycero-3-phosphocholine, cholesterol, and a polyethylene glycol-lipid^50^. This mixture was rapidly combined with an aqueous solution containing mRNA at pH 4. The ionizable cationic lipid (pKa in the range of 6.0–6.5), proprietary to Acuitas Therapeutics (Vancouver, Canada), and LNP composition are described in the United State patent application WO2017/004143A1. The average hydrodynamic diameter was 80 nm with a polydispersity index of 0.02–0.06 as measured by dynamic light scattering using a Zetasizer Nano ZS (Malvern Instruments, Malvern, UK) and an encapsulation efficiency of 95% as determined using a Quant-iT Ribogreen assay (Life Technologies). Encapsulated mRNAs were aliquoted at 30 μl per tube (1 mg/ml) for single usage and stored at −80°C. The empty LNP (eLNP) without encapsulated mRNAs were similarly purified and used as negative controls for vaccines.

### LNP vaccine treatment

For neoantigen vaccine treatment, 10 µg of LNP vaccine was mixed in 50 µl of PBS and injected subcutaneously on Day 10 or Day 17 post-inoculation.

### Immunohistochemistry

Tumor tissues were washed with cold PBS and fixed in 4% paraformaldehyde (PFA), paraffin-embedded, and sectioned onto slides for immunohistochemistry (IHC). The slides were deparaffinized with xylene, rehydrated, and antigen retrieved in a microwave oven for 20 min. After the inhibition of endogenous peroxidase activity, the slides were incubated overnight at 4°C with anti-mouse CD3 antibody (Abcam) and anti-mouse BHLHE40 antibody. The slides were then incubated with EnVision reagent (Dako, Glostrup, Denmark), and a color reaction was developed using 2% 3,3-diaminobenzidine in 50 mM tris buffer (pH 7.6) containing 0.3% hydrogen peroxidase. Coverslips were mounted on glass slides, and imaged using a ##Evos FL Auto Imaging System. Post-processing of the images was done using a Fiji v2.14.0 software.

### Mouse Primary T cell isolation and activation

The mouse primary CD8^+^ T cells were isolated from mouse spleens by a positive selection method with a mouse CD8a + T Cell isolation kit, following the supplier’s protocol. Mouse primary T cells were activated using a mouse T Cell Activation/Expansion Kit, and cultured in RPMI-1640 supplemented with 10% FBS, 1% HEPES, 1% P/S, under 37 °C in a 5% CO_2_ incubator.

### Tumor cell isolation

Tumor tissues thoroughly minced into fragments smaller than 1 mm and suspended in type IV collagenase solution (1.0–1.5 mg ml^−1^). The tumor samples with digestion medium were incubated for 1 h at 37 °C and 5% CO_2_ incubator and then passed through a 70 µm filter to obtain a single-cell suspension. The cell suspension was washed with RPMI-1640 media by centrifuging at 350 × g for 5 min (twice). The resulting single-cell tumor suspension was then used for FACS analysis. Cells (10^6^ cells per 100 µl) were labelled for CD3 using antibodies (Invitrogen, 1:100) and neoantigen tetramer (a gift from NIH Tetramer Core Facility^32^;1:300) for 90 min on ice. The cells were washed twice with PBS, and analyzed using an Aurora CS2 Cell Sorter Spectral Flow Cytometer. Mouse cancer cells were isolated from mice tumor by a negative selection method using anti-mouse CD3 antibody, and then purified by a negative selection method using mouse CD45+ Cell isolation kit. Tumor cell isolation efficiency varied among samples, and in some cases, tumor-infiltrating T cells or cancer cells could not be successfully recovered in sufficient numbers. Only samples with adequate viable cell yields were included in downstream analyses. Final sample sizes are reported in the corresponding figure legends.

### Cell mitochondria labeling and coculture

Cells were stained with 0.5–1.0 µM of MitoTracker Green FM (MG) or MitoTracker Deep Red FM (MR) at 37 °C in a 5% CO_2_ incubator as per the manufacturer’s protocols. The MitoTracker-stained cells were washed twice, followed by incubation in complete media for 1 h to allow removal of excess unbound MitoTracker dyes. Cocultures were established by incubating cancer cells and immune cells in a 5:1 ratio of IL-2-free RPMI-1640 media for 3 hours under 37 °C in a 5% CO_2_ incubator. For anti-human CD38 antibody treatment, A549 cells were pretreated with 10µg/ml Isatuximab antibody for 24 h prior to coculture.

### Flow cytometry

Cell suspensions were prepared in PBS containing 2% FBS at a density of 1–5 million cells ml^−1^. Depending on the experimental design, fixation, permeabilization and staining were performed as stated above. Anti-human CD8 antibody, anti-mouse CD8 antibody, and anti-mouse CD3 antibody (Invitrogen) were used at 1:50 dilution for flow cytometry analysis. Samples were analyzed using a CytoFlex S (6 lasters) Flow Cytometer or an Aurora CS2 Cell Sorter Spectral Flow Cytometer. All the FACS data were processed with FlowJo v10.5.3 software. For flow sorting, a single-cell suspension was prepared from coculture or tumor tissues; the cells were sorted according to their fluorescence signal. The cells were collected in a 5 ml flow tube containing FBS, transferred to fresh medium, and counted before subsequent experiments.

### Tunneling nanotube labeling

Cells grown on coverslips (VWR; diameter, 12 mm) were fixed with 4% paraformaldehyde at room temperature for 2 h. The fixed cells were washed three times with 1× PBS for 20 min. Cells were then permeabilized with 0.5% Triton X-100 at 4 °C for 10 min. Next, the cells were washed three times with 1× PBS-T (1× PBS + 0.05% Tween 20), blocked with 10% bovine serum albumin (BSA) at room temperature for 1 h, and stained with the 50 μg ml^−1^ of phalloidin AF 405 for 1 h at room temperature. The cells were washed three additional times with 1× PBS-T. Coverslips were mounted on glass slides and imaged using a Zeiss LSM 980 Confocal with ZEN Blue v3.5 software. Post-processing of the images was done using a Fiji v2.14.0 software.

### RNA isolation and quantitative real-time PCR (real-time qPCR)

Total RNA was extracted by using a TRIzol reagent according to the manufacturer’s protocol. The concentration of total RNA in each sample was measured using a NanoDrop 2000 Spectrophotometer. For mRNA detection, equal amounts of RNA was reverse transcribed into cDNA using High Capacity cDNA Reverse Transcription Kit. Real-time qPCR was performed on a QuantStudio 5 using Power SYBR Green PCR Mater Mix. mRNA expression levels were normalized to the endogenous control GAPDH. Relative mRNA expressions were quantified by 2^−ΔΔCt^. Primers are listed in **Table S5**. Real-time qPCR reactions were performed using the following cycling conditions: initial denaturation at 95 °C for 10 min, followed by 40 cycles of amplification at 95 °C for 15 s and 60 °C for 1 min.

### Fluorescence in situ hybridization and quantification of human mitochondrial signals in mouse tumors

Mice were injected subcutaneously in both flanks with wild-type MC38 cells (1 × 10 cells per flank). Beginning on day 14, a total of 1 × 10 human primary T cells (wild-type ‘WT’, BHLHE40 overexpression ‘BHLHE40 OE’, S100A4 knockdown ‘S100A4 KD’, or combined ‘OE+KD’) were injected subcutaneously between the two tumors on the dorsal side of each mouse every two days. Tumors were harvested on day 32, washed with cold phosphate-buffered saline (PBS), fixed in 4% paraformaldehyde (PFA), paraffin-embedded, and sectioned onto glass slides. Fluorescence in situ hybridization (FISH) was performed using RNAscope assay obtained from Advanced Cell Diagnostics (ACD) according to the manufacturer’s protocol. The RNAscope probe design algorithm by ACD examined each oligo probe for potential cross-reactivity against the host transcriptome to ensure the probe specificity. There was a verification procedure following each major step during probe design to guarantee the accuracy. Probe quality was verified by ACD through *in silico* verifications. Probe design algorithm selected oligos complementary to the input target sequence with compatible melting temperature for optimal hybridization at RNAscope assay condition and minimal cross-hybridization to off-target sequences. During the manufacture and formulation process, oligo synthesis QC was performed on each oligo probe. The oligo synthesis QC included ESI mass spec to make sure the probe sequences were not containing any deletion, addition or mutations, and OD reading to ensure correct concentration was provided upon delivery.

For this study, two RNAscope probes were synthesized: 1) human CD3E to identify T cells, and 2) a custom human-specific mitochondrial probe (Hs-MT-No-XMm), to specifically detect human mtDNA. CD3E probe was visualized with ClariTSA Fluorophore 570 to identify T cells, and Hs-MT-No-XMm was visualized using ClariTSA Fluorophore 520.

Sections were counterstained with DAPI to visualize cell nuclei. Coverslips were mounted on glass slides and imaged using a Zeiss LSM 980 Confocal with ZEN Blue v3.5 software. Post-processing of the images was done using a Fiji v2.14.0 software. To assess the presence of human mitochondrial signals in neighboring mouse tumor cells, six microscopic fields were selected from each tumor section. Within each field, five human primary T cells were selected as reference cells. Human T cells were defined as cells containing a DAPI-positive nucleus associated with a human CD3E signal. Fluorophore 570 signals that were not spatially associated with a DAPI-positive nucleus were considered nonspecific and excluded from the analysis. DAPI-positive, human CD3E-negative cells located within a 50-μm radius of each selected human T cell were evaluated as candidate mouse tumor cells. Cells lacking an identifiable DAPI-positive nucleus were excluded. A candidate mouse tumor cell was classified as positive for human mtDNA if one or more discrete Hs-MT-No-XMm/Fluorophore 520 puncta were located within the boundaries of an intact DAPI-defined cell. Fluorophore 520 signals located outside intact cells, lacking spatial association with a DAPI-positive cell, or otherwise considered inconsistent with intracellular punctate staining were excluded as nonspecific signals. The number of Hs-MT-No-XMm-positive candidate mouse tumor cells within the 50-μm radius was subsequently quantified and compared across groups.

### Absolute human mitochondrial DNA copy number quantification

Mice were subcutaneously injected in the flanks with wild-type MC38 cells (1 × 10 cells per injection). Starting on day 14, a total of 1 × 10 human primary T cells (wild-type ‘WT’, BHLHE40 overexpression ‘BHLHE40 OE’, S100A4 knockdown ‘S100A4 KD’, or combined ‘OE+KD’) were injected subcutaneously between the two tumors on the dorsal side of each mouse every two days. Tumors were harvested on day 32. Single-cell suspensions were prepared, and human CD8 T cells, mouse CD8 T cells, and mouse CD45 tumor cells were isolated using a Human CD8a+ T Cell Isolation Kit, a Mouse CD8a+ T Cell Isolation Kit, and a Mouse CD45+ Cell Isolation Kit, respectively, according to the manufacturer’s instructions. Total DNA was extracted using the DNeasy Blood & Tissue Kit. Human mitochondrial DNA (mtDNA) copy number in mouse tumor cells was quantified using the Absolute Human Mitochondrial DNA Copy Number Quantification qPCR Assay Kit following the manufacturer’s protocol.

### Western blotting

Cells were seeded at a density of 2–3 × 10^6^ cells in T-25 flasks. Cells were lysed with radioimmunoprecipitation assay (RIPA) lysis buffer (supplemented with protease and phosphatase inhibitor) for 30 min on ice with gentle vortexing every 10 min, followed by centrifugation at 4 °C. Protein concentration was measured using a bicinchoninic acid (BCA) assay according to the supplier’s protocol. Equal amounts of protein lysates were electrophoresed on a 10–15% polyacrylamide gel at 80–150 V for 1 h. The proteins were then transferred to polyvinylidene difluoride (PVDF) membrane (0.2 µm pore size) by wet transfer for 2 h at 220 mA (Bio-Rad Laboratories). Membranes were blocked with 5% BSA in 1× TBS-T (1× TBS and 0.1% Tween 20) for 1 h and incubated with primary antibodies at 4 °C for 12–14 h. After washing thrice with 1× TBS-T, membrane was incubated with the secondary HRP-conjugated anti-mouse or anti-rabbit antibodies (1:1000) for 1 h at room temperature. Membranes were then washed twice with 1X TBS-T and once with TBS before developed using chemiluminescent Femto substrate. Images were acquired using an Azure Biosystems imaging system and analyzed with Sapphire v1.2.1228.0 software. The CD38 antibody (1:500) were used for Western blotting.

### Determination of CD38/ADPR-cyclase activity

Nicotinamide guanine dinucleotide (NGD^+^) was used to determine the ADPR-cyclase activity by fluorometric analysis. For human ADPR-cyclase activity analysis, the CD38 KO A549 cells were transduced with lentivirus expressing human wild-type CD38 or the E226D mutation for 24–48 h, followed by selection of transduced cells using puromycin (0.5–10 μg/mL). For mouse ADPR-cyclase activity analysis, the Cd38 KD MC38 cells were transduced with lentivirus expressing mouse wildtype Cd38 or E230D mutation for 24–48 h, followed by puromycin selection (0.5–10 μg/mL). Cells were then incubated with 200 μM NGD^+^, and the production of cyclic GDP-ribose (cGDPR) was measured at excitation/emission wavelengths of 300 nm/ 410 nm using a SYNERGY H1 microplate reader with Gen5 v3.17 software.

### IFN-γ enzyme-linked immunospot (ELISpot) assay

Human cancer cells and human primary T cells were cocultured in a 5:1 ratio of IL-2-free RPMI-1640 medium for 3 h under 37 °C in a 5% CO_2_ incubator. The culture supernatant was then collected, and IFN-γ levels were measured using a human IFN-γ ELIS kit according to the manufacturer’s protocol. Mouse primary T cells were sorted using an Aurora CS2 Cell Sorter Spectral Flow Cytometer, and the cell numbers were determined using a Countess 3 automated cell counter with trypan blue staining. The cells were then cultured IL-2-free RPMI-1640 medium for 3 h under 37 °C in a 5% CO_2_ incubator. Culture supernatants were collected, and IFN-γ levels were measured using a mouse IFN-γ ELISA kit according to the manufacturer’s protocol.

### Reactive Oxygen Species (ROS) levels assessment

DCFDA/ H2DCFDA-Cellular ROS Assay Kit was used to measure ROS levels in cell cultures according to the manufacturer’s protocol. Cells were harvested and prepared as a single-cell suspension as described above. The cells were then stained with 20 μM DCFDA in culture medium and incubated for 30 minutes at 37 °C. After incubation, cells were washed once with 1× Buffer and suspended in medium. Cell numbers were determined using a Countess 3 automated cell counter, and samples were immediately analyzed at excitation/emission wavelengths of 488/535 nm using aSYNERGY H1 microplate reader with Gen5 v3.17 software.

### ATP Production assessment

Total cellular ATP levels were measured using the ATP Determination Kit according to the manufacturer’s protocol. Cells were harvested and prepared as a single-cell suspension as described above, and the cell numbers were determined using a Countess 3 automated cell counter with trypan blue staining. Cells were lysed with lysis buffer (10 mM Tris (pH7.5), 100 mM NaCl, 1 mM EDTA, 0.01% Triton X-100) for 10 min on ice. Lysates were then diluted 1:10 with the ATP Determination Kit reaction mix to a final volume of 200 μl and immediately analyzed at an emission wavelength of 560 nm using a SYNERGY H1 microplate reader with Gen5 v3.17 software.

### Cell apoptosis assay

Cell apoptosis was measured using an Annexin V-FITC Apoptosis Detection Kit according to the manufacturer’s protocol. Briefly, cells were harvested and prepared as a single-cell suspension as described above, and cell numbers were determined using a Countess 3 automated cell counter with trypan blue staining. Cells were resuspended in binding buffer at a concentration of 1 × 10 cells/mL. Then, 5 µL of Annexin V-FITC conjugate and 10 µL of propidium iodide (PI) solution were added to each cell suspension. After incubation, apoptosis levels were analyzed using a CytoFLEX S (6 lasers) flow cytometer or an Aurora CS2 Spectral Flow Cytometer. All FACS data were processed using FlowJo v10.5.3 software.

### Cell viability assay

WT or CD38 KD A549 cells and MC38 cells (5 × 10^5^ cells per well) were cultured in 6-well plates under normal conditions. Cells were harvested at 1 h, 2 h, 3 h, and 4 h using trypsin and counted using a Countess 3 automated cell counter with trypan blue staining. All experiments were performed in triplicate, and results are expressed as total number of cells in each well.

### Single-cell RNA-seq library preparation and sequencing

Tumor cells from WT and KD mice were harvested, prepared as a single-cell suspension as described above, and stained with anti-mouse CD3 antibody (Invitrogen) at a 1:50 dilution. Samples were sorted using an Aurora CS2 Spectral Flow Cytometer. Subsequently, 10,000 CD3^+^ and CD3^-^ cells were separately loaded onto the Chromium Controller (10X Genomics, PN-120223) using a Chromium Single Cell Chip K (10X Genomics, PN-1000286) and processed to generate single-cell gel beads in emulsion (GEMs) according to the manufacturer’s protocol (10X Genomics, CG000331). Libraries were generated using Chromium Single Cell 5’ Reagent Kits v2 (10X Genomics, PN-1000244) and TCR Amlification Kit (10X Genomics, PN-1000254) according to the manufacturer’s instructions. Library quality control was performed using an Agilent TapeStation 4200 (G2991A) with High Sensitivity D5000 reagents. The libraries were sequenced on NovaSeq X Plus (PE150-10B with approximately 375 Gb of theoretical data per lane.

### Single-cell RNA-seq data analysis

The Cell Ranger v7.1.0 Software Suite was used to process the raw scRNA-seq data into gene expression profiles. FASTQ files for each sample were processed with cellranger count or multi using mm39 reference genome. Scanpy was used to analyze the gene count matrices^51^. Low-quality cells were filtered out if the number of expressed genes was less than 200 or the percentage of mitochondrial transcripts was greater than 50%. We used a loose mitochondrial transcript percentage threshold because the percentage could reflect real biology in the context of mitochondrial transfer. Despite this loose threshold, most cells had a mitochondrial transcript percentage of less than 5%. Genes were filtered out if they were detected in less than 3 cells. Scrubletwas used to detect doublets^52^. The gene count matrix was log-normalized and the top 2,000 highly variable genes were selected. Principal component analysis (PCA) was performed on the log-normalized data using the 2,000 highly variable genes. Harmonywas used to remove batch effects in PCA space^53^. Cd3 expression was defined as the minimum UMI count among Cd3d, Cd3e, and Cd3g and cells with Cd3 expression > 0 were defined as T cells and used for subsequent analysis.

UMAP was performed on the batch-corrected neighborhood graph to visualize T cells on a two-dimensional layout. The Leiden algorithm was used to cluster T cells into subgroups at a resolution of 1.0, producing a total of 12 clusters. Differentially expressed genes were identified using the Wilcoxon test after excluding mitochondrial, ribosomal, and hemoglobin genes. Four clusters were removed because they expressed markers of both T cells and another cell type and were therefore likely doublets. After the filtering steps, 8 clusters corresponding to 17,181 T cells were retained for downstream analyses.

We defined CD4 T cells as cells with Cd4 expression but not Cd8a or Cd8b1 expression, CD8 T cells as cells with Cd8a or Cd8b1 expression but not Cd4 expression, double-positive T cells as cells expressing both markers, and double-negative T cells as cells expressing neither. We then defined the lineage of each T cell cluster: mixed if both CD4 and CD8 T-cell percentages were greater than 30%, and CD4 or CD8 if one lineage was greater than 40%. We defined cell types based on marker gene expression.

Differentially expressed genes between WT and KD groups were identified using the Wilcoxon test after excluding ribosomal, mitochondrial, and hemoglobin genes. Gene set enrichment analysis (GSEA) was then performed on the gene list ranked by Wilcoxon scores using GSEApy^54^. The pathway gene sets used were extracted from Molecular Signatures Database curated mouse canonical pathway collection, including pathways from BioCarta, Reactome, and WikiPathways^55^. Pathways were considered significant if Benjamini–Hochberg adjusted FDR was < 0.05.

### BHLHE40 ChIP-seq analysis

BHLHE40 ChIP-seq alignment BAM files for GM12878 were downloaded from ENCODE4^46^: ENCFF827XJM and ENCFF314PKF. H3K4me3 ChIP-seq alignment BAM files for human T cells were downloaded from ENCODE4: ENCFF167PUH and ENCFF822TCP^48^. BHLHE40 ChIP-seq raw read files for Jurkat cells were downloaded from GEO: GSE122940^47^ and aligned using Bowtie2. ENCODE3 candidate cis-regulatory elements (cCREs) were downloaded from the UCSC Genome Browser. S100A4 isoform information was obtained from GENCODE v45. pyGenomeTracks was used to generate genome browser tracks.

### Kaplan-Meier estimates

The patients were downloaded from Timer2^56^ and stratified into two groups (BHLHE40 high and low) with cutpoints selected by maximizing log-rank statistic. LogHR was evaluated using cox regression with age as covariate. Statistical significance was evaluated using log rank test. LGG, Brain Lower Grade Glioma. PAAD, Pancreatic adenocarcinoma. GBM, Glioblastoma multiforme. LUSC, Lung squamous cell carcinoma.

### Statistical analysis

Data are expressed as mean ± s.e.m. Statistical analyses were performed using Prism GraphPad 9.5.0 software. Each data point represents an individual animal. No data points were excluded from the analyses. Data were analyzed using one-way ANOVA followed by Tukey’s post hoc test or a two-sided Wilcoxon rank-sum test. A *p* value < 0.05 was considered statistically significant.

## Supplemental Figure Legends

**Figure S1. Multiple human and mouse cancer cell lines acquire mitochondria from T cells.**

(A) Representative images of all cancer cell lines and human primary T cells used in this study.

(B) Schematic of the flow cytometry analysis strategy.

(C) Flow cytometry analysis of MitoTracker Deep Red (MR) fluorescence intensity in U2OS cancer cells cultured alone or co-cultured with human primary T cells. **** *P* < 0.0001. Results are the mean ± SE (n = 3).

(D) Flow cytometry analysis of MitoTracker Deep Red (MR) fluorescence intensity in PC9 cancer cells cultured alone or co-cultured with human primary T cells. **** *P* < 0.0001. Results are the mean ± SE (n = 3).

(E) Flow cytometry analysis of MitoTracker Deep Red (MR) fluorescence intensity in MC38 cancer cells cultured alone or co-cultured with mouse primary T cells. **** *P* < 0.0001. Results are the mean ± SE (n = 3).

(F) Flow cytometry analysis of MitoTracker Deep Red (MR) fluorescence intensity in B16/F10 cancer cells cultured alone or co-cultured with mouse primary T cells. **** *P* < 0.0001. Results are the mean ± SE (n = 3).

**Figure S2. Inhibition of intercellular mitochondrial transfer.**

(A) Schematic illustration of the transwell coculture system of T cells and A549 cells, followed by flow cytometric analysis of mitochondrial transfer in A549 cells.

(B) Quantification of mitochondrial transfer shown in (A). * *P* < 0.05, *** *P* < 0.001, **** *P* < 0.0001. Results are the mean ± SE (n = 3).

(C) Effect of shRNA-mediated knockdown on apoptosis in A549 cancer cells, measured by flow cytometry (FITC percentage).

(D) Effect of TLN2 knockdown on mitochondrial transfer, measured by flow cytometry (MitoTracker Deep Red percentage). ** *P* < 0.01. Results are the mean ± SE (n = 3).

(E) Effect of TLN2 knockdown on apoptosis in A549 cancer cells, measured by flow cytometry (FITC percentage). *** *P* < 0.001. Results are the mean ± SE (n = 3).

(F) Flow cytometry analysis of mitochondrial transfer and apoptosis in WT and TLN2 KD A549 cells. Results are the mean ± SE (n = 3).

(G) Correlation analysis between gene knockdown and apoptosis in WT and TLN2 KD A549 cells.

(H) Effects of shRNA-mediated double knockdown in A549 cells. * *P* < 0.05, ** *P* < 0.01. Results are the mean ± SE (n = 3).

(I) Mitochondrial transfer in WT and KD A549 cancer cells following co-culture with T cells, measured by flow cytometry (MitoTracker Deep Red percentage). This experiment represents an independent repeat of Figure 1E. *** *P* < 0.001, **** *P* < 0.0001. Results are the mean ± SE (n = 6).

(J) Effect of cytarabine (Ara-C) treatment on apoptosis in A549 cancer cells, measured by flow cytometry (FITC percentage). Results are the mean ± SE (n = 3).

(K) CD38 expression in tumor and normal tissues analyzed using the GEPIA platform.

**Figure S3. CD38 regulates TNT formation through its ligand-binding function rather than its NAD-catalytic activity.**

(A) mRNA expression of CD38 in MC38 cells following CRISPR–Cas9-mediated CD38 knockout and reconstitution with WT or E230D mutant CD38. Results are the mean ± SE (n = 6).

(B) Flow cytometry analysis of the proportion of WT and E230D MC38 cancer cells acquiring T cell–derived mitochondria following 3 h co-culture.

(C) Quantification of mitochondrial transfer shown in (B). Results are the mean ± SE (n = 6).

(D) Bulk RNA expression of CD38 across tissues analyzed using the GTEx platform.

(E) Bulk RNA expression of CD31 (PECAM1) across tissues analyzed using the GTEx platform.

(F) Single-cell RNA expression of CD38 across tissues analyzed using the GTEx platform.

(G) Single-cell RNA expression of CD31 (PECAM1) across tissues analyzed using the GTEx platform.

(H) Flow cytometry analysis of mitochondrial acquisition by WT A549 cells (stained with MitoTracker Green, MG) following co-culture with WT, CD38 KD, or CD31 KD A549 cells (stained with MitoTracker Deep Red, MDR). * *P* < 0.05, ** *P* < 0.01, *** *P* < 0.001. Results are the mean ± SE (n = 3).

Figure S4. CD38 knockdown markedly impairs tumor growth *in vivo*.

(A) Schematic of subcutaneous implantation of WT and CD38 KD MC38 tumor cells in mice.

(B) mRNA expression of Cd38 in tumor-derived CD45 cells. ** *P* < 0.01. Results are the mean ± SE (n = 6 tumors for WT, n = 3 tumors for KD).

(C) Body weights of tumor-bearing mice. Results are the mean ± SE (n = 5).

(D) Tumor growth curves of WT and CD38 KD MC38 tumors. ** *P* < 0.01, *** *P* < 0.001, **** *P* < 0.0001. Results are the mean ± SE (n = 5).

(E) Tumors size at Day 23.

(F) Schematic of the Tet-On shRNA system.

(G) mRNA expression of Cd38 in MC38 cells under the indicated conditions: WT, Tet-On shRNA, and Tet-On shRNA with doxycycline (1 μg/mL, 24 h). * *P* < 0.05. Results are the mean ± SE (n = 3).

(H) mRNA expression of Cd38 in tumor-derived CD45 cells following Tet-On shRNA induction. ** *P* < 0.01. Results are the mean ± SE (n = 3).

(I) Cell proliferation of A549 cells with or without CD38 knockdown. Results are the mean ± SE (n = 3).

(J) Cell proliferation of MC38 cells with or without CD38 knockdown. Results are the mean ± SE (n = 3).

**Figure S5. Expression of Bhlhe40 and S100a4 in scRNA-seq data and their functional effects in T cells.**

(A) Violin plot showing median gene count, median UMI count, and mitochondrial reads (%) in the scRNA-seq dataset.

(B) Marker genes defining T cell subtypes identified in the scRNA-seq dataset.

(C) Volcano plots showing differentially expressed genes (DEGs) in CD8-Eff and Mixed-Act clusters from KD versus WT tumors.

(D) Dot plot showing the correlation between Bhlhe40 and S100a4 expression in T cells from scRNA-seq data.

(E) Effect of transwell on mitochondrial transfer between T cells and A549 cells. *** *P* < 0.001, **** *P* < 0.0001. Results are the mean ± SE (n = 3).

(F) Reactive oxygen species (ROS) levels in human T cells under the indicated conditions: wild-type (WT), BHLHE40 overexpression (BHLHE40 OE), S100A4 knockdown (S100A4 KD), and combined BHLHE40 overexpression with S100A4 knockdown (OE+KD). * *P* < 0.05, ** *P* < 0.01. Results are the mean ± SE (n = 6).

(G) ATP levels in human T cells under the indicated conditions as in (D). ** *P* < 0.01, *** *P* < 0.001. Results are the mean ± SE (n = 6).

(H) IFN-γ (IFNG) production in human T cells under the indicated conditions as in (D). ** *P* < 0.01. Results are the mean ± SE (n = 6).

**Figure S6. The cross-species conservation of mitochondria transfer**

(A) Fluorescence in situ hybridization of human mitochondrial signals in mouse tumors sections from Figure 4A. In the merged images, regions containing positive human mitochondrial signals are outlined with red dashed lines. The punctate green signals represent positive signals detected by the human mitochondrial probe, whereas the diffuse green staining represents nonspecific background signals. CD3E-positive signals within the outlined regions are indicated by yellow arrows in the Human CD3E+ panels. Cell nuclei are visualized by DAPI staining. The specificity of the human mitochondrial signals was further evaluated by examining the presence or absence of CD3E-positive cells and DAPI signals within the corresponding regions, which helped distinguish true positive human mitochondrial signals from nonspecific background signals. Scale bars, 50 μm.

(B) Human mtDNA copy number measured by a qPCR-based assay in A549 cells, MC38 cells, and a blank control (n = 3).

(C) Protein sequence alignment of human S100A4 and mouse S100a4 generated using ClustalW.

**Figure S7. Expression and functional effects of CD31, BHLHE40, and S100A4 in naïve and activated T cells.**

(A) mRNA expression of CD31 (PECAM1) in naïve and activated human T cells. ** *P* < 0.01, **** *P* < 0.0001. Results are the mean ± SE (n = 6).

(B) Flow cytometry analysis of mitochondrial transfer to A549 cancer cells co-cultured with naïve or activated T cells under the indicated conditions (WT, BHLHE40 OE, S100A4 KD, and OE+KD).

(C) Quantification of mitochondrial transfer shown in (D). * *P* < 0.05, *** *P* < 0.001, **** *P* < 0.0001. Results are the mean ± SE (n = 3).

(D) Quantification of CD8 T cell numbers in (D). ** *P* < 0.01, *** *P* < 0.001. Results are the mean ± SE (n = 3).

(E) mRNA expression of BHLHE40 in naïve and activated human T cells. * *P* < 0.05, *** *P* < 0.001, **** *P* < 0.0001. Results are the mean ± SE (n = 6).

(F) mRNA expression of S100A4 in naïve and activated human T cells. * *P* < 0.05, ** *P* < 0.01, **** *P* < 0.0001. Results are the mean ± SE (n = 6).

**Figure S8. Antigen-specific (Tetramer□) and non-antigen-specific (Tetramer□) T cells in tumors**

(A) Flow cytometry analysis of sorting strategy for tetramer□ and tetramer□ T cells from tumors shown in Figure 5A.

(B) Reactive oxygen species (ROS) levels in tetramer□ and tetramer□ T cells isolated from tumors in Figure 5A. * *P* < 0.05, ** *P* < 0.01. Results are the mean ± SE (n = 6 tumors for Ctrl-WT and Ctrl-KD; n = 8 tumors for LNP-WT; n = 9 tumors for LNP-KD).

(C) mRNA expression of Cd31 in tetramer□ and tetramer□ T cells isolated from tumors in Figure 5A. * *P* < 0.05. Results are the mean ± SE (n = 6 tumors for Ctrl-WT and Ctrl-KD; n = 8 tumors for LNP-WT; n = 9 tumors for LNP-KD).

(D) ATP levels in tetramer□ and tetramer□ T cells isolated from tumors in Figure 5A. * *P* < 0.05, ** *P* < 0.01. Results are the mean ± SE (n = 6 tumors for Ctrl-WT and Ctrl-KD; n = 8 tumors for LNP-WT; n = 9 tumors for LNP-KD).

(E) IFN-γ (IFNG) production in tetramer□ and tetramer□ T cells isolated from tumors in Figure 5A. * *P* < 0.05. Results are the mean ± SE (n = 6 tumors for Ctrl-WT and Ctrl-KD; n = 8 tumors for LNP-WT; n = 9 tumors for LNP-KD).

---

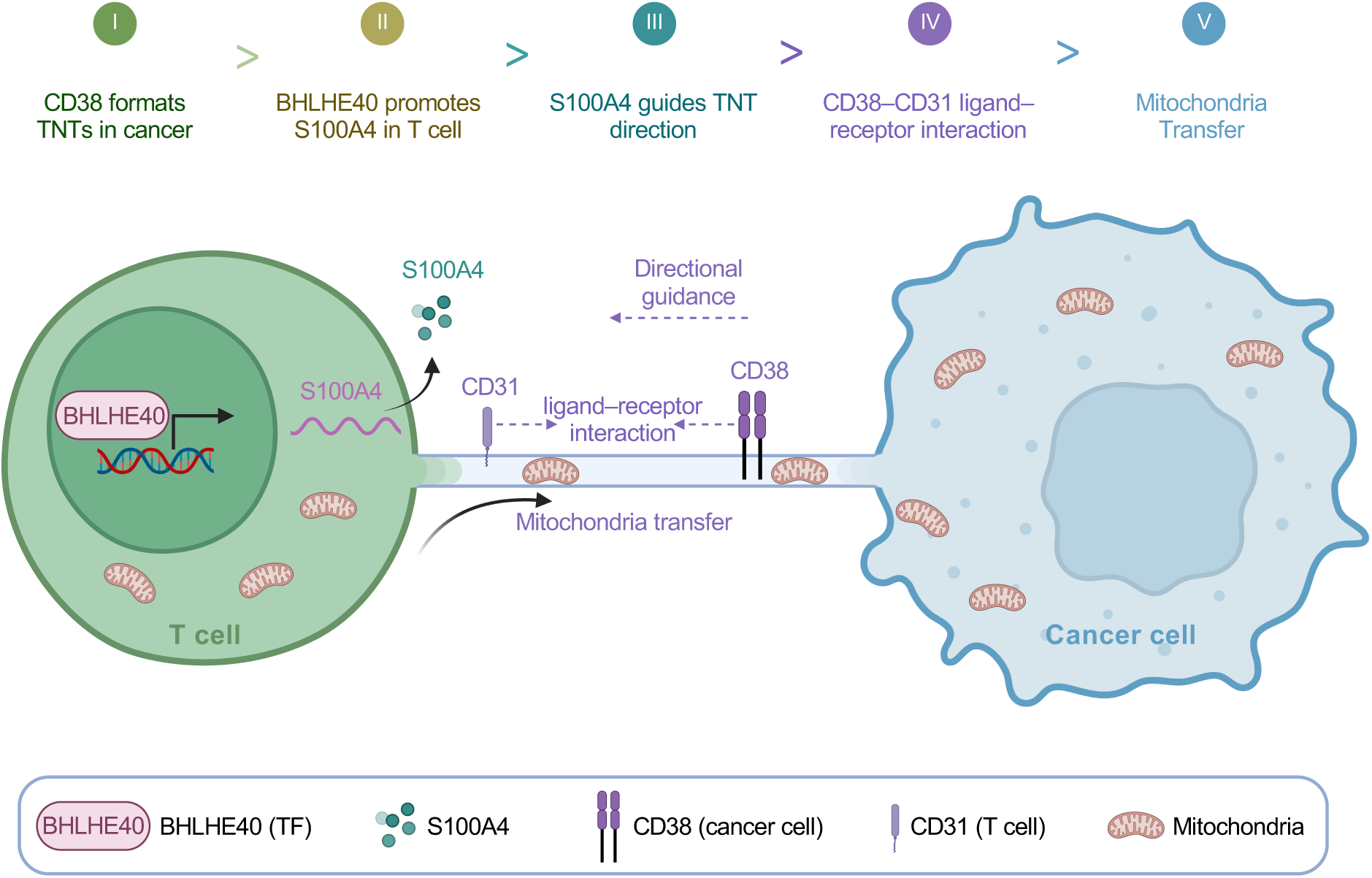

