## Supplementary figures and images for "Cancer cells hijack mitochondria from Bhlhe40+ T cells for immune evasion"

### Supplemental Figure 1

Figure S1

A

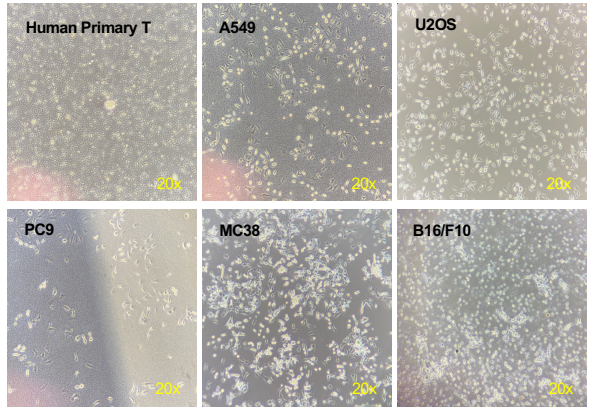

B

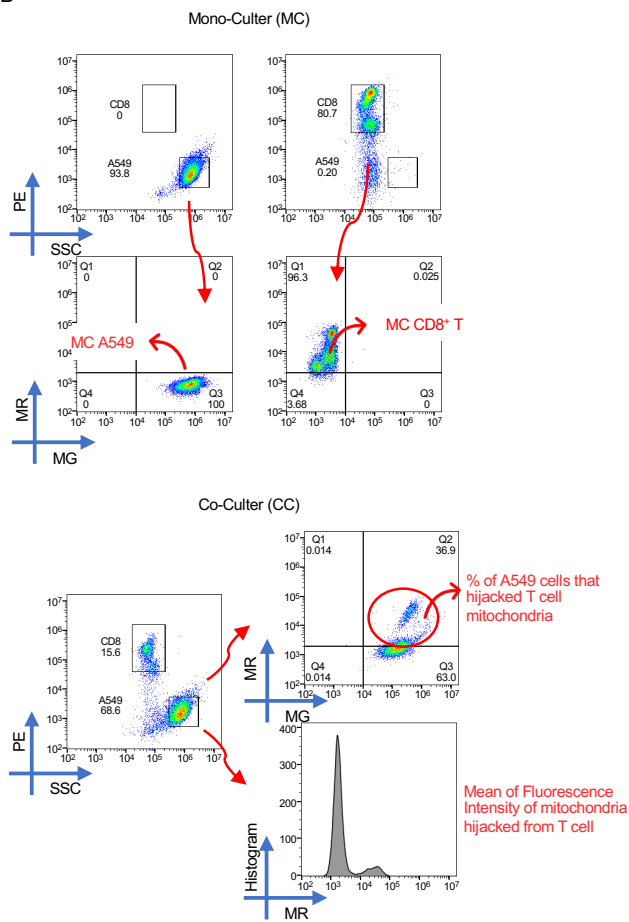

C

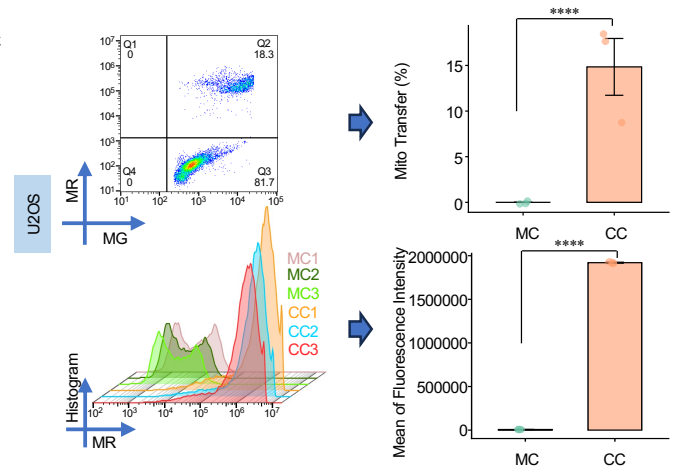

D

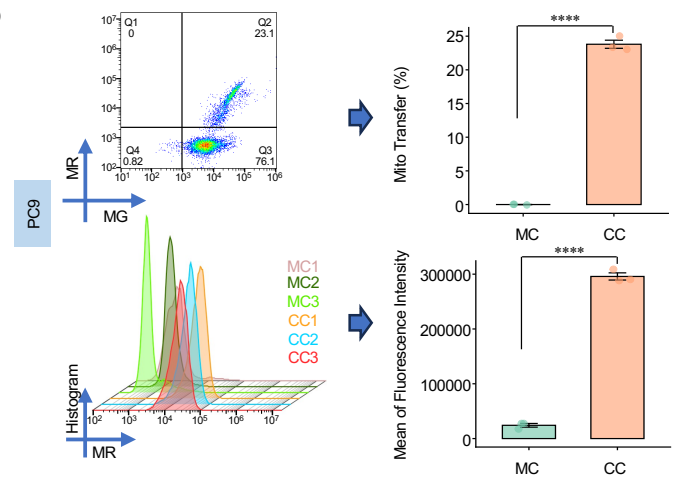

E

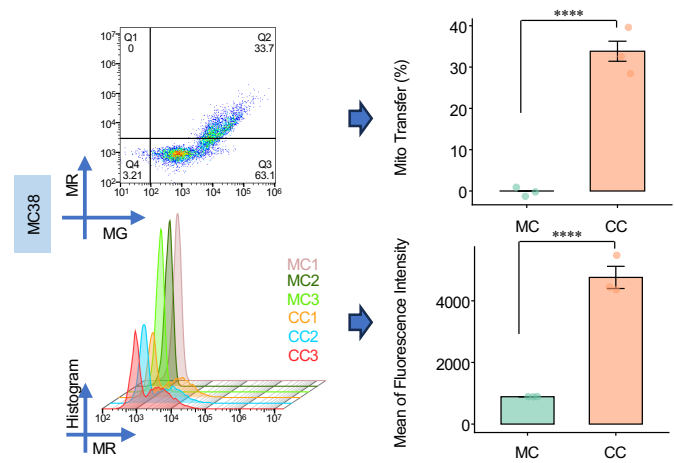

F

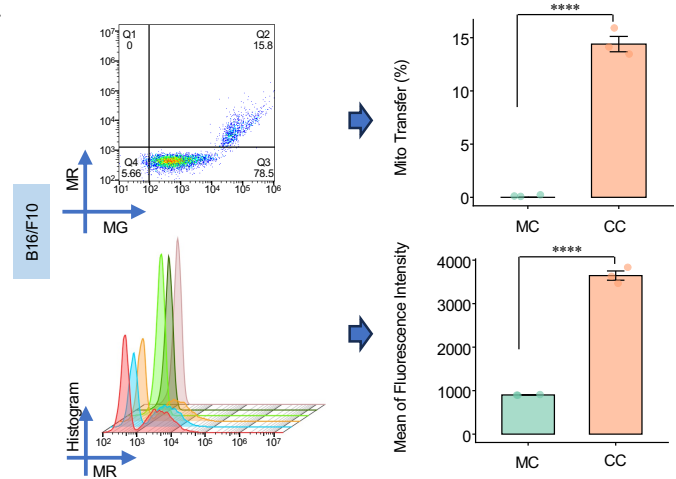

### Supplemental Figure 2

Figure S2

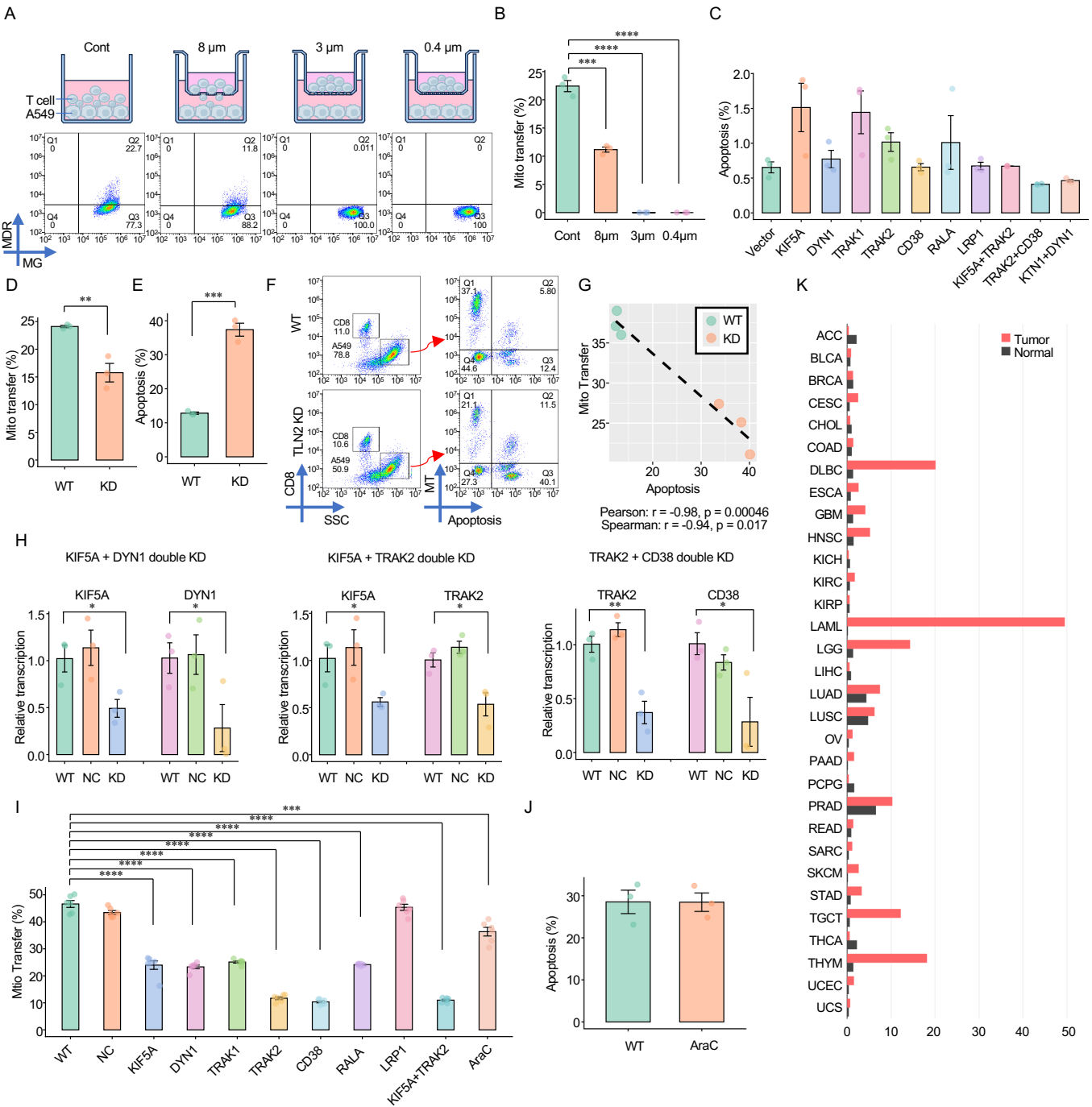

### Supplemental Figure 3

Figure S3

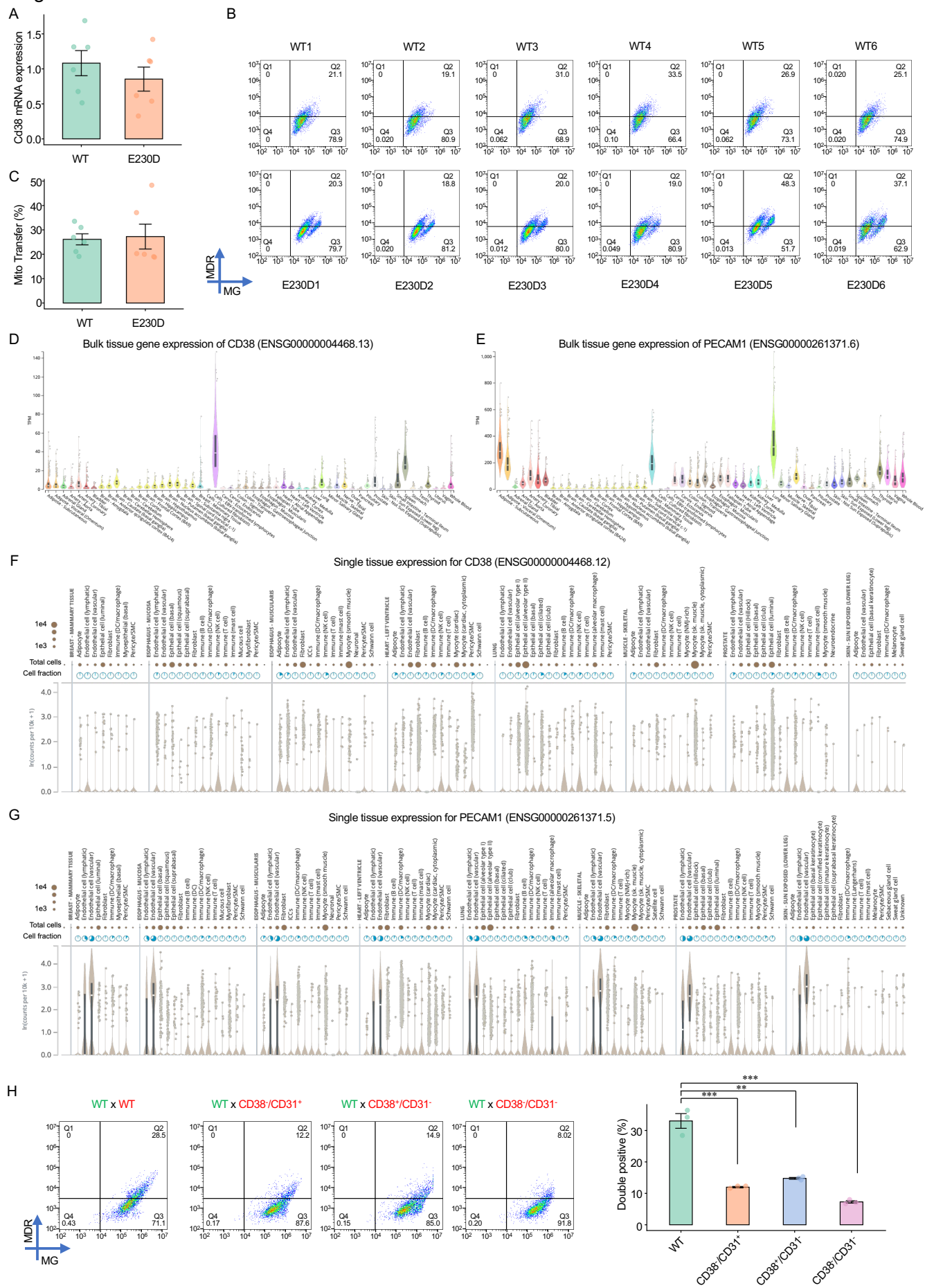

### Supplemental Figure 4

Figure S4

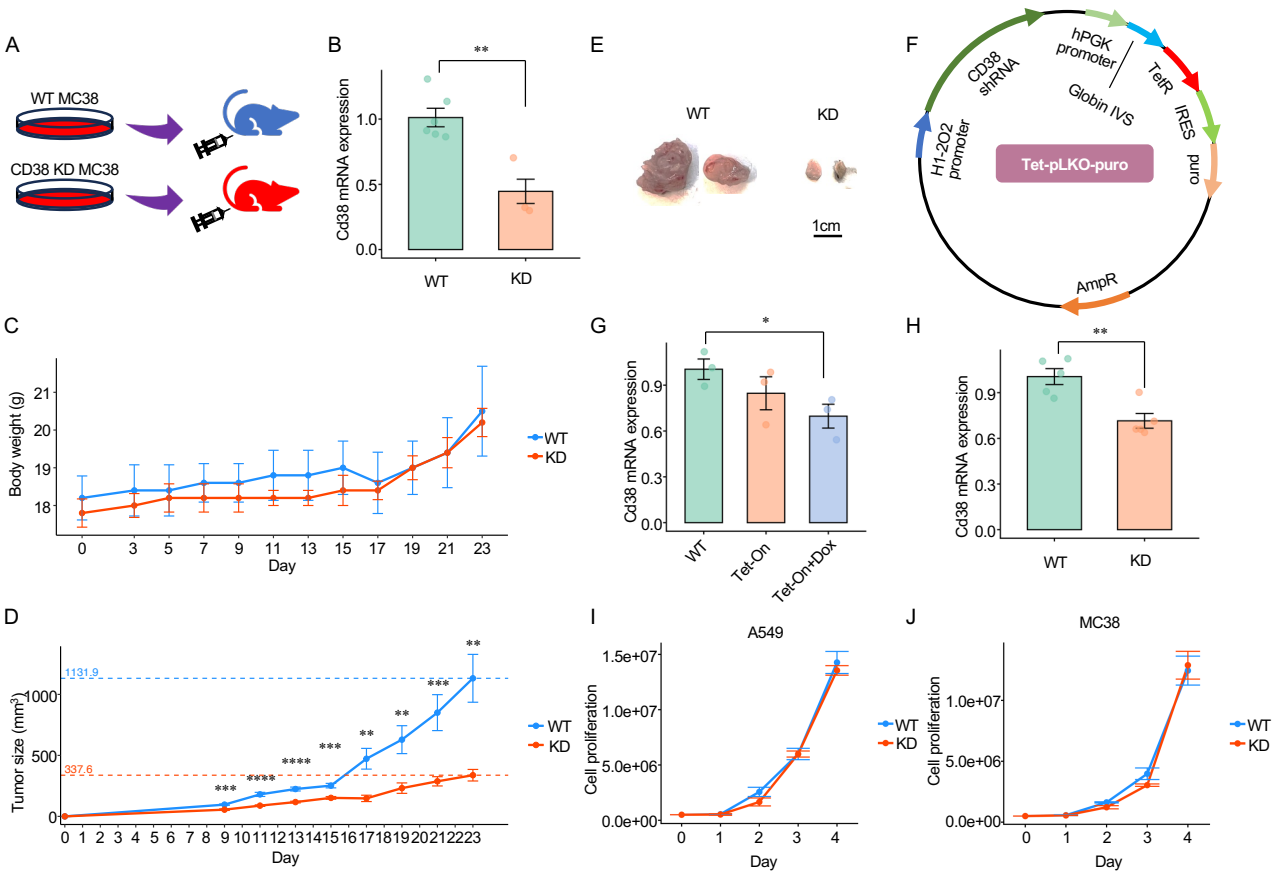

### Supplemental Figure 5

Figure S5

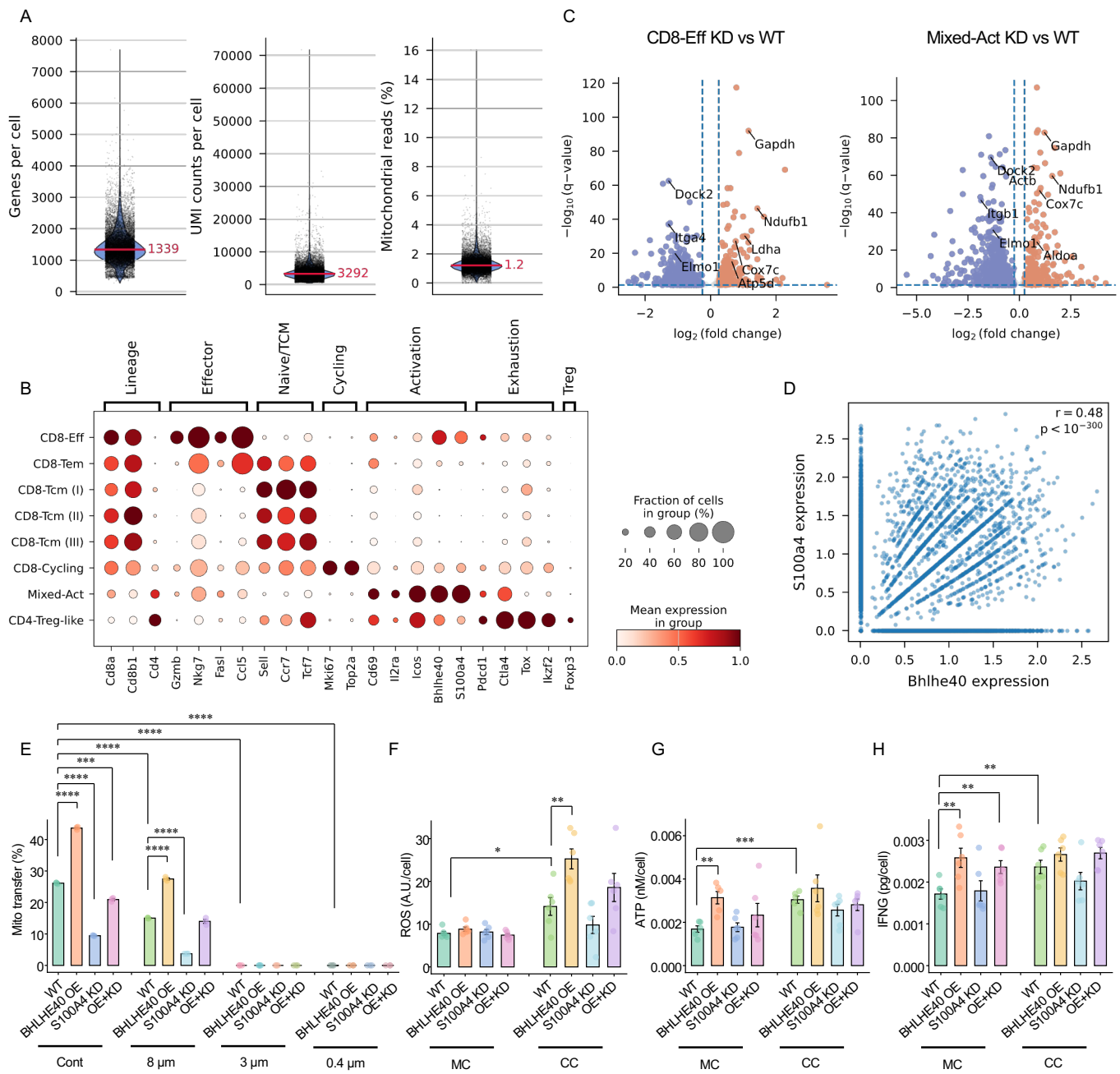

### Supplemental Figure 6

Figure S6

A

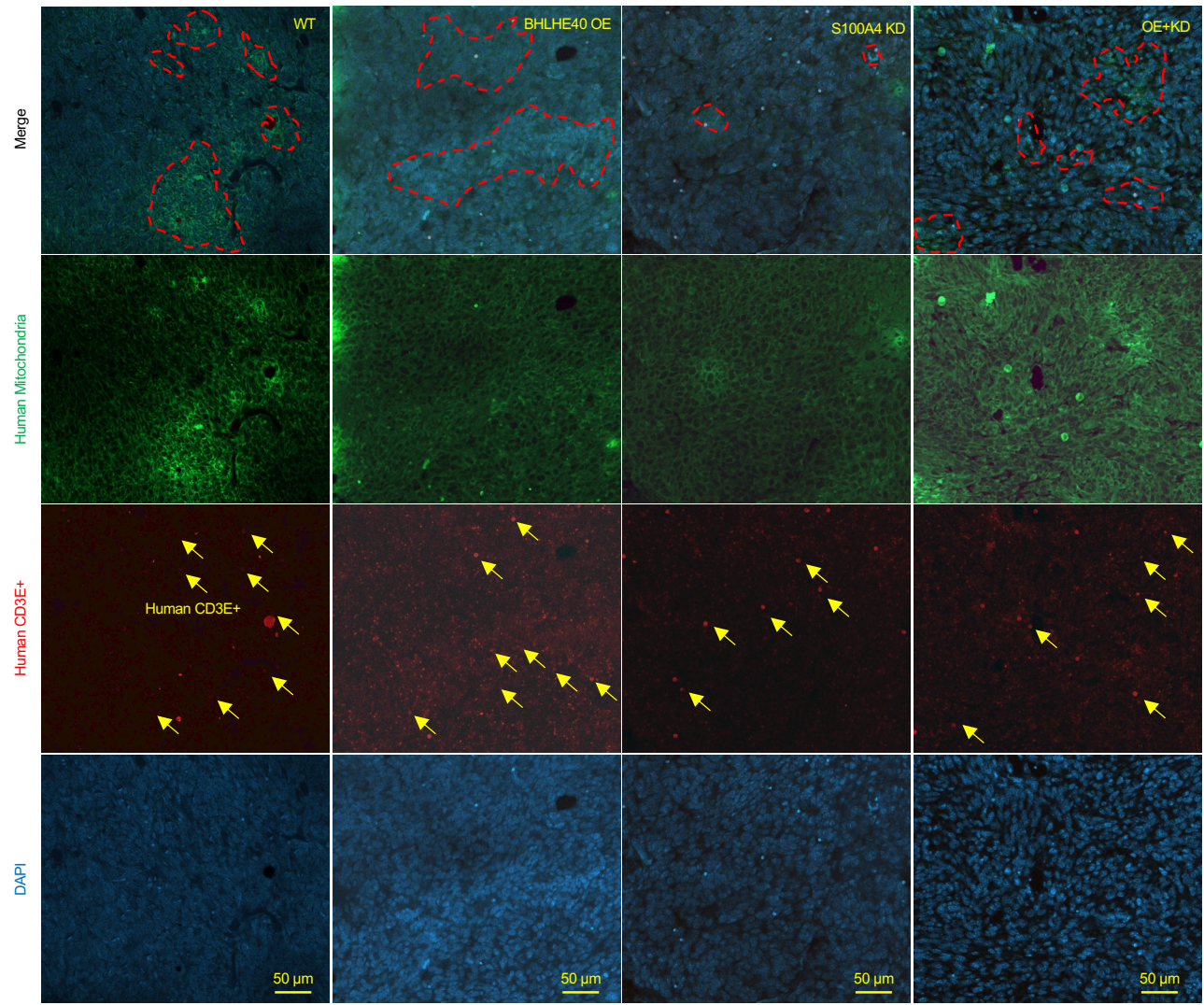

B

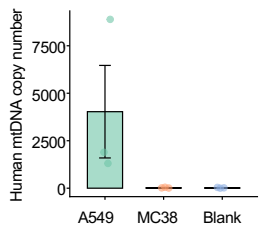

C

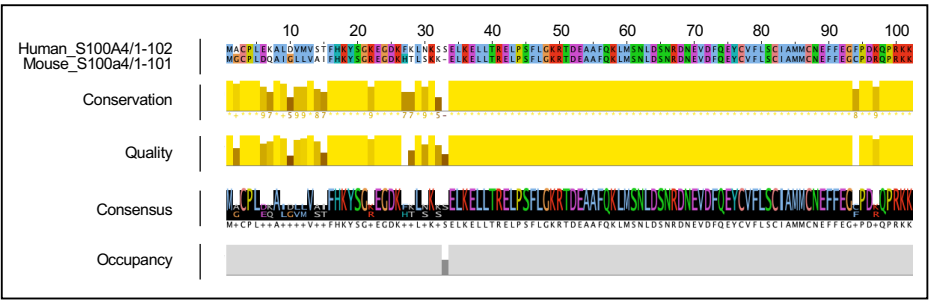

### Supplemental Figure 7

Figure S7

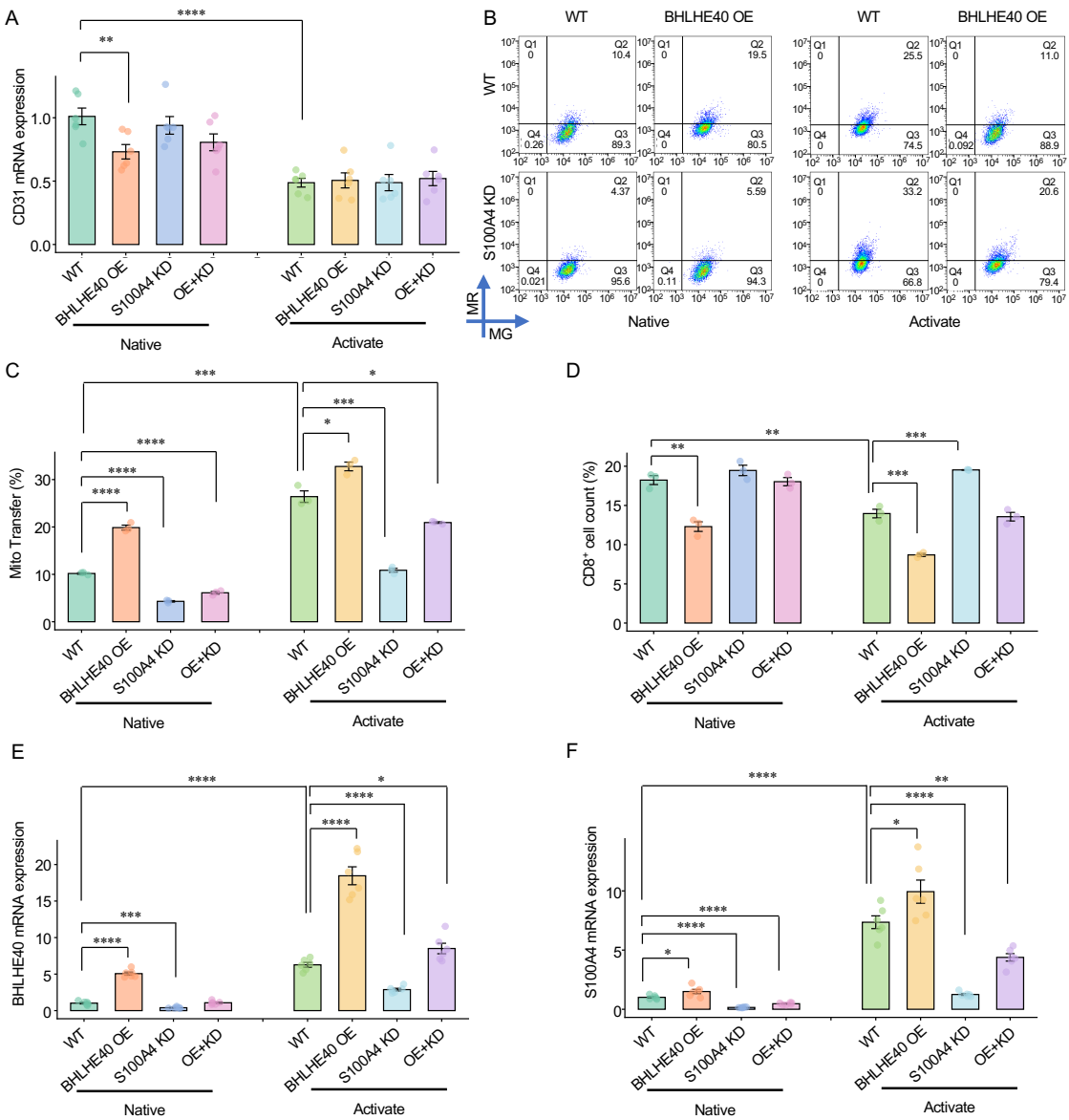

### Supplemental Figure 8

Figure S8

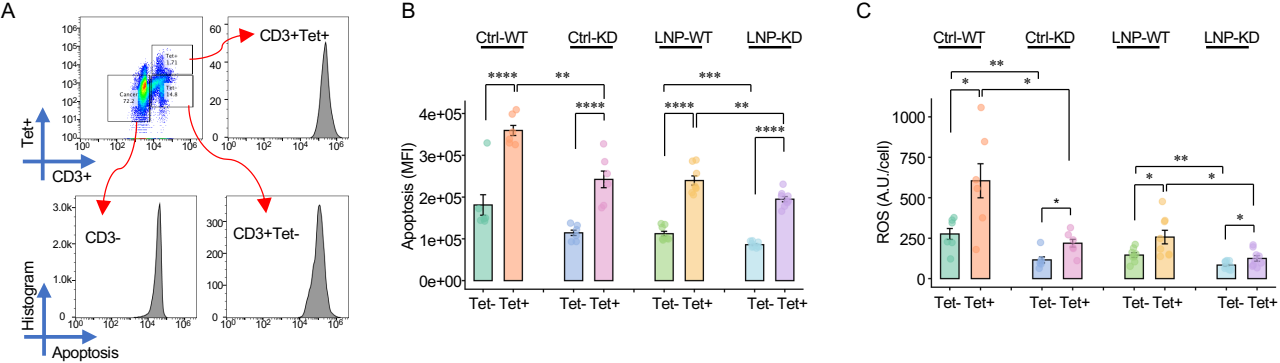
